# Mapping immune cellular landscapes and vaccine responses across a spectrum of health and immunodeficiency

**DOI:** 10.64898/2026.05.21.726432

**Authors:** Davide Vespasiani, Annelise Quig, James Lancaster, Cindy Yuting Shen, Jack Cooper, Zewen Kelvin Tuong, Amanda Jackson, Susanne Schulz, Shuk-Yin (Sylvia) Tsang, Kirsten Deckert, Erin C Lucas, Mai Margetts, Miles Horton, Samantha Chan, Julian J Bosco, Josh Chatelier, Samar Ojaimi, Charlotte Slade, Celina Jin, Hamish W King

**Affiliations:** Walter and Eliza Hall Institute of Medical Research, VIC Australia; Department of Medical Biology, University of Melbourne, VIC Australia; Ian Frazer Centre for Children’s Immunotherapy Research, University of Queensland, QLD Australia; Department of Clinical Immunology and Allergy, Royal Melbourne Hospital, VIC Australia; Department of Infectious Diseases & Immunology, Austin Health, VIC Australia; Asthma, Allergy and Clinical immunology, Alfred Hospital, VIC Australia; School of Translational Medicine, Monash University, VIC Australia; Department of Pathology, Monash Health, VIC Australia; Department of Medicine, Southern Clinical School, Monash University, VIC Australia; Department of Pathology, Royal Melbourne Hospital, VIC Australia; Department of Infectious Diseases, University of Melbourne, VIC Australia

## Abstract

Immune responses to infection and vaccination exhibit diversity between individuals that can be shaped by differences in their immune cell landscapes and the signalling, transcriptional, and genetic mechanisms that coordinate immune cell function. Specific antibody deficiency (SAD) and common variable immunodeficiency (CVID) are common forms of predominantly antibody deficiencies that result in poor responses to vaccination. While molecular and cellular causes of the immune dysfunction and poor vaccination responses for individuals with CVID have been reported, immune cell or molecular defects have not yet been identified in SAD. Here, we have used single-cell multi-omics to define the cellular landscapes, transcriptional states, adaptive immune repertoires and protein expression of patients with SAD and CVID before and after polysaccharide vaccination. We discovered that while SAD and CVID exhibit overlapping immune defects, including accumulation of exhausted NK memory cells and dysregulated expression of genes that mediate lipopolysaccharide sensing and clearance by monocytes, individuals with SAD have a unique expansion of cytotoxic CD4^+^ T cells that correlates with reduced regulatory T cells. In response to vaccination, we observed rapid changes in gene expression associated with lipopolysaccharide responses by monocytes and NF-kB pathway activation in B cells, and an apparent expansion of a CD95^+^ class-switched memory B cell population that does not occur in patients with lower antigen-specific responses. Together, our findings reveal cellular and molecular factors that underpin variability in vaccine responses and define SAD in a broader spectrum of immune dysfunction.

## INTRODUCTION

Inter-individual variability in vaccine responses is well established, with some individuals mounting robust, durable immunity while others show reduced or transient protection^1–5^. Early studies have focused on demographic and clinical factors such as age, sex, prior antigen exposure, and comorbidities, which accounts for part of this variation^1^. Different antigen types also elicit distinct responses; protein antigens typically induce T cell-dependent germinal-centre (GC) derived responses, whereas polysaccharide-based vaccines are thought to drive T cell-independent responses^6^. It has been of considerable interest to investigate the diversity of cellular and molecular responses to vaccination by different antigen types^2,3,5^. Predominantly antibody deficiencies (PADs) are the most common form of inborn errors of immunity, a heterogeneous group of disorders caused by genetic defects that compromise immune function and regulation^7–9^. The investigation of PADs has advanced our understanding of normal immune function and development, as exemplified by the identification of Bruton’s tyrosine kinase deficiency as a cause of X-linked agammaglobulinaemia^10^. As such, investigating the cellular composition, transcriptional states, and signalling pathways that shape vaccine responses across different PADs provides a clinically relevant framework to define how these mechanisms are disrupted and to compare immune dysfunction across disease states.

PADs clinically present with recurrent sinopulmonary bacterial infections and are characterised by reduced immunoglobulin levels or impaired antibody responses to specific antigens. Despite the significant morbidity associated with these diseases, arising from complications of infection and accrual of permanent end-organ damage, delayed diagnosis is a common problem^11^. Two of the most common PADs associated with diagnostic challenges are specific antibody deficiency (SAD) and common variable immunodeficiency (CVID), where diagnosis for the latter relies upon criteria such as reduced immunoglobulin levels, absent isohemagglutinins and/or poor vaccine responses^12,13^. In contrast, SAD patients have normal immunoglobulin levels but impaired antibody responses to polysaccharide antigens, with measurement of antibody responses to the 23-valent pneumococcal polysaccharide vaccine traditionally used to diagnose patients. However, use of different assays to measure antibody titres and lack of standardised definition for ‘normal’ polysaccharide vaccine responses has complicated the classification of SAD^14,15^, even though it is a leading indication for immunoglobulin replacement therapy worldwide^16,17^.

The reduced ability of patients with SAD to mount responses to polysaccharide antigens suggests a defect in T-independent immune pathways, but so far limited flow cytometry-based studies have only identified modest reductions in class-switched memory B cells^18^ and increases of immature CD5^+^ B cells^19^. The cellular and molecular basis of SAD therefore remains unclear, making it difficult to place within the broader spectrum of clinical immune dysfunction. This includes how it relates to conditions such as CVID, which often has clear immune cell defects such as reductions in class-switched memory B cells and plasma cells^8,20^, altered CD4^+^ T cell^21^ and NK cell^22^ phenotypes and higher frequencies of classical and intermediate monocytes^23^. Alternatively, SAD could reflect a lower end of phenotypic variation in responses to polysaccharide-based vaccines without underlying immune dysfunction.

We therefore set out to investigate and define the spectrum of immune responses to a typhoid Vi-polysaccharide antigen across a cohort encompassing SAD, CVID and healthy donors. We leveraged single-cell multi-omics to generate a comprehensive resource characterizing shared immune dysfunction between PADs including dysregulated expression of genes that mediate lipopolysaccharide sensing and clearance, and accumulation of exhausted NK cells. Intriguingly, we identify a SAD-specific expansion of cytotoxic CD4⁺ T cells that may be associated with reduced regulatory T cell states in this cohort. We describe vaccination-induced transcriptional responses in monocytes and B cells, and expansion of class-switched memory B cells, which were absent in individuals with poor antigen-specific responses. Together, our findings define cellular and molecular features underlying variability in vaccine responses and place SAD within a broader spectrum of adaptive immune dysfunction for the first time.

## RESULTS

### A single-cell multi-omic atlas defines dysfunctional immune cell landscape of SAD

To address the current lack of mechanistic understanding of SAD and its position within a spectrum of immune dysfunction, we generated a single-cell multi-omic atlas of peripheral immune cells from patients with SAD and CVID **(Figure 1a, Table S1**). All patients with SAD and CVID were on immunoglobulin replacement therapy and over the 13-month study period experienced more infections requiring oral antibiotic treatment compared with healthy donors (**Figure S1a**). With the exception of one CVID individual who was heterozygous for a pathogenic allele in *TRNT1* (p.Ser418fs) and another harboring two variants of unknown significance in *TTC7A* (p.Lys606Arg, p.Ser672Pro) previously observed in CVID^24^, all other CVID and SAD participants were not found to carry known pathogenic or likely pathogenic variants in genes associated with inborn errors of immunity. We performed single-cell cellular indexing of transcriptomes and epitopes by sequencing (CITE-seq) to profile gene expression, cell surface protein expression and B and T cell receptor (B/TCR) sequences in peripheral blood mononuclear cells (PBMCs) (**Figure 1a**). After demultiplexing and quality control, we retained 165,641 single-cells (median 5,823 cells per donor) and annotated 22 major cell type clusters covering all expected major immune cell lineages (**Figure 1b, Figure S1b-g**). We identified broad differences in cell type frequencies for most major immune cell lineages in SAD and CVID compared with healthy controls (**Figure 1c-d, Table S2**), consistent with immune dysfunction in both cohorts. Both SAD and CVID had reduced plasma cell frequencies, which was less pronounced for SAD and consistent with typically milder immunoglobulin defects associated with this condition^14,25–27^. An increased abundance of unswitched memory B cells was observed only in CVID (**Figure 1e**)^8,20^. Both cohorts exhibited increased classical CD14^+^ monocytes and CD56^low^ NK cells (**Figure 1d, Figure S1h**), suggesting a common inflammatory landscape in these two diseases. Evidence for unique immune dysfunction in SAD was an enrichment of effector memory CD8^+^ T cells (**Figure S1g**).

**Figure 1.**
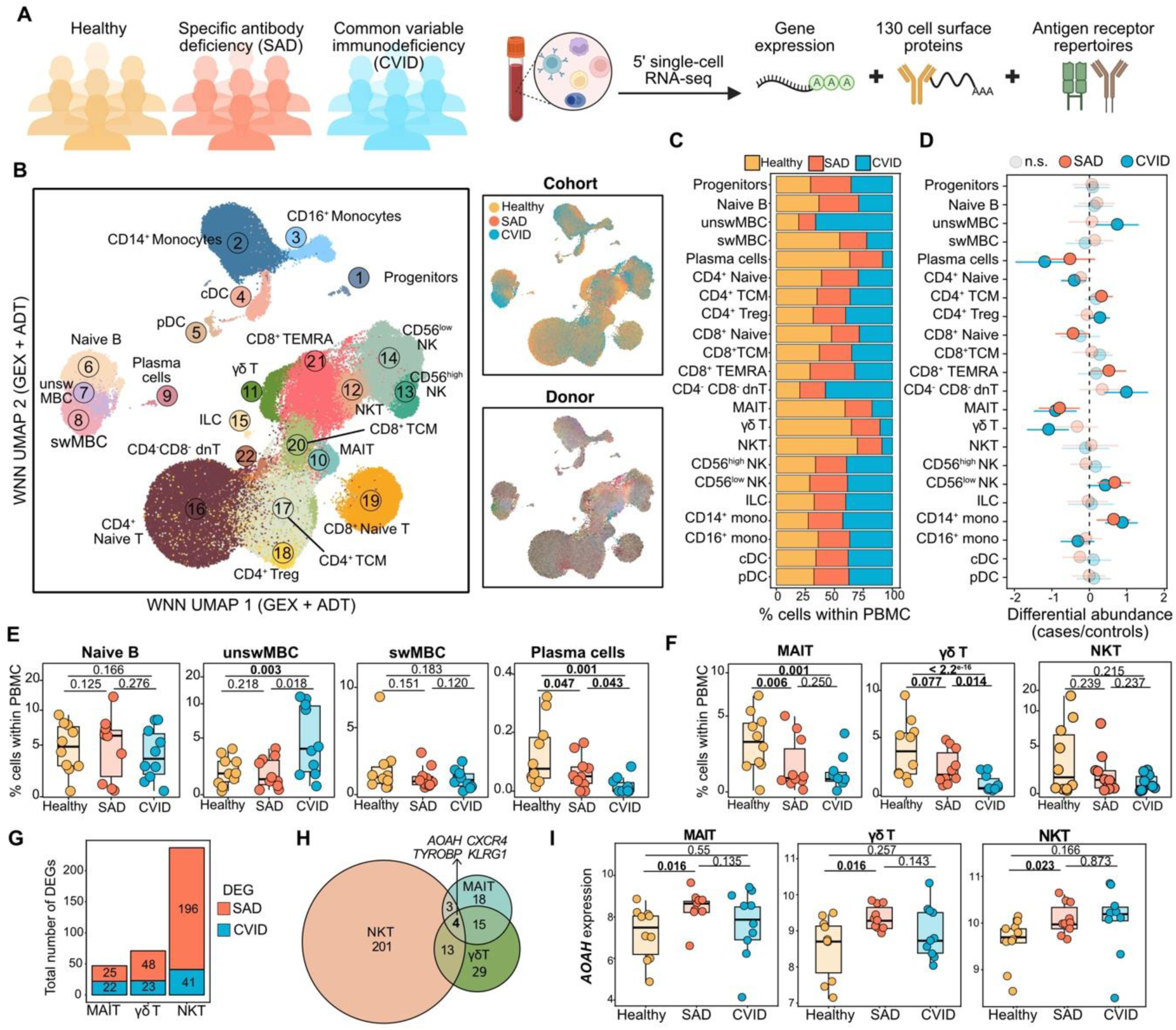
A single-cell atlas of SAD and CVID identifies shared and unique immune dysfunction. **A)** Schematic of PAD cohorts and single-cell atlas generation. **B)** Weighted nearest neighbour Uniform Manifold Approximation and Projection (WNN UMAP) of CITE-seq data for 165,641 high-quality single-cells colored by cell type annotation, cohort or donor. **C)** Cell type frequencies across cohorts. **D)** Relative cell type frequencies between cases and controls. Error bars represent 95% credible interval; dashed black bars represent fold difference of 0.1. Opaque dots and bars indicate significant results as per sccomp FDR-corrected *p*-value. **E)** Cell type frequency relative to total PBMC of B cell subsets. *p* values as per D. **F)** Cell type frequency relative to total PBMC of non-conventional T cell subsets. **G)** Total number of differentially expressed genes (DEG) identified between cases and controls within non-conventional T cells **H)** Overlapping DEGs between three non-conventional T cell types. **I)** Pseudobulk expression (log2 counts per million) of *AOAH* across cohorts within MAIT (left), γδ T (middle) and NKT cells (right). *p* denotes Student’s t-test result.

Reductions in non-conventional T cells (mucosal-associated invariant T (MAIT), γδ T and natural killer T (NKT) cells) were also observed for both SAD and CVID^28–30^, albeit with some patient-to-patient variability (**Figure 1f**). Non-conventional T cells bridge innate and adaptive immunity by recognising non-peptide antigens, mounting rapid effector responses, and contributing to inflammatory signalling. To understand potential functional differences in these cells, we compared gene expression between cohorts and found more differentially expressed genes (DEGs) in SAD for all non-conventional T cell populations examined (**Figure 1g**). Of the four genes that were differentially expressed in all three cell types (**Figure 1h**), *AOAH* was more highly expressed in SAD (**Figure 1i**). *AOAH* encodes acyloxyacyl hydrolase which is an evolutionary conserved host lipase involved in the inactivation of lipopolysaccharide (LPS)^31^. Together, our single-cell multi-omic resource reveals that SAD is associated with distinct alterations in both adaptive and innate immune cell populations and that impaired innate sensing may contribute to increased susceptibility to bacterial infections.

### Dysfunctional CD14^+^ monocyte states are features of SAD and CVID

CD14⁺ monocytes are key mediators of innate responses to microbial polysaccharides, facilitating pathogen recognition and inflammatory signalling through pathways such as CD14 and TLR4. In CVID, elevated plasma levels of soluble CD14 and CD14^+^ myeloid cells have previously been associated with defective responses to LPS^23,32,33^, however, it remains unclear if defective signalling by CD14^+^ monocytes or other myeloid cells occurs in SAD^14,25,26^. Although we found evidence for reduced expression of activation-related genes in CD16^+^ monocytes (**Figure S2a-b**), most differential gene expression between PAD and healthy donors were observed in CD14^+^ monocytes (**Figure S2a**). This included elevated expression of inflammatory and interferon-related genes (*IFI44L*, *OAS3*, *IFI6, STAT1, IRF1;* **Figure 2a-b**), suggesting that CVID and SAD have a shared inflammatory phenotype. Given altered expression of LPS-related genes in non-conventional T cells, we were intrigued to identify an enrichment for genes associated with cellular responses to LPS, including LPS sensing (*TLR4, LY86*^34^); LPS-mediated inflammatory responses (*CD86, LITAF*^35,36^) and clearing of LPS from macrophage cell surface (*ABAC1*^37^; **Figure 2b-c**). Compared with healthy controls, there was lower expression of these LPS-related genes suggesting potential dysregulation of responses to LPS by CD14^+^ monocytes in SAD. To investigate whether these differences originated from a specific CD14^+^ monocyte state, we further clustered myeloid cells to identify nine transcriptionally distinct cell states, including five CD14^+^ monocytes (basal, activated, IFN^high^, *EREG*^high^, *G0S2*^high^); CD14^+^CD16^+^ intermediate monocytes; CD16^+^ monocytes and four dendritic cell subsets (*AXL*^+^ *SIGLEC6*^+^ DC, cDC1, cDC2 and pDC) (**Figure 2d-e, Figure S2c-d**). In keeping with elevated expression of IFN-related genes in CD14^+^ monocytes, we identified significant enrichment of IFN^high^ CD14^+^ monocytes in CVID (**Figure 2f-g**) defined by high expression of genes associated with interferon responses (**Figure 2h**), suggesting this monocyte state may be a source of a reported interferon signature in CVID^32,38^. We also observed reduced frequencies of activated HLA-DR^+^CD14⁺ monocytes (**Figure 2i-j**) and this activated population was noted to have elevated expression of genes involved in LPS responses (**Figure 2k**, **Figure S2d**). This suggests defects in CD14⁺ monocyte activation contribute to impaired LPS sensing and downstream immune responses in both CVID and SAD which may affect responses to bacterial infections and polysaccharide-based vaccines.

**Figure 2.**
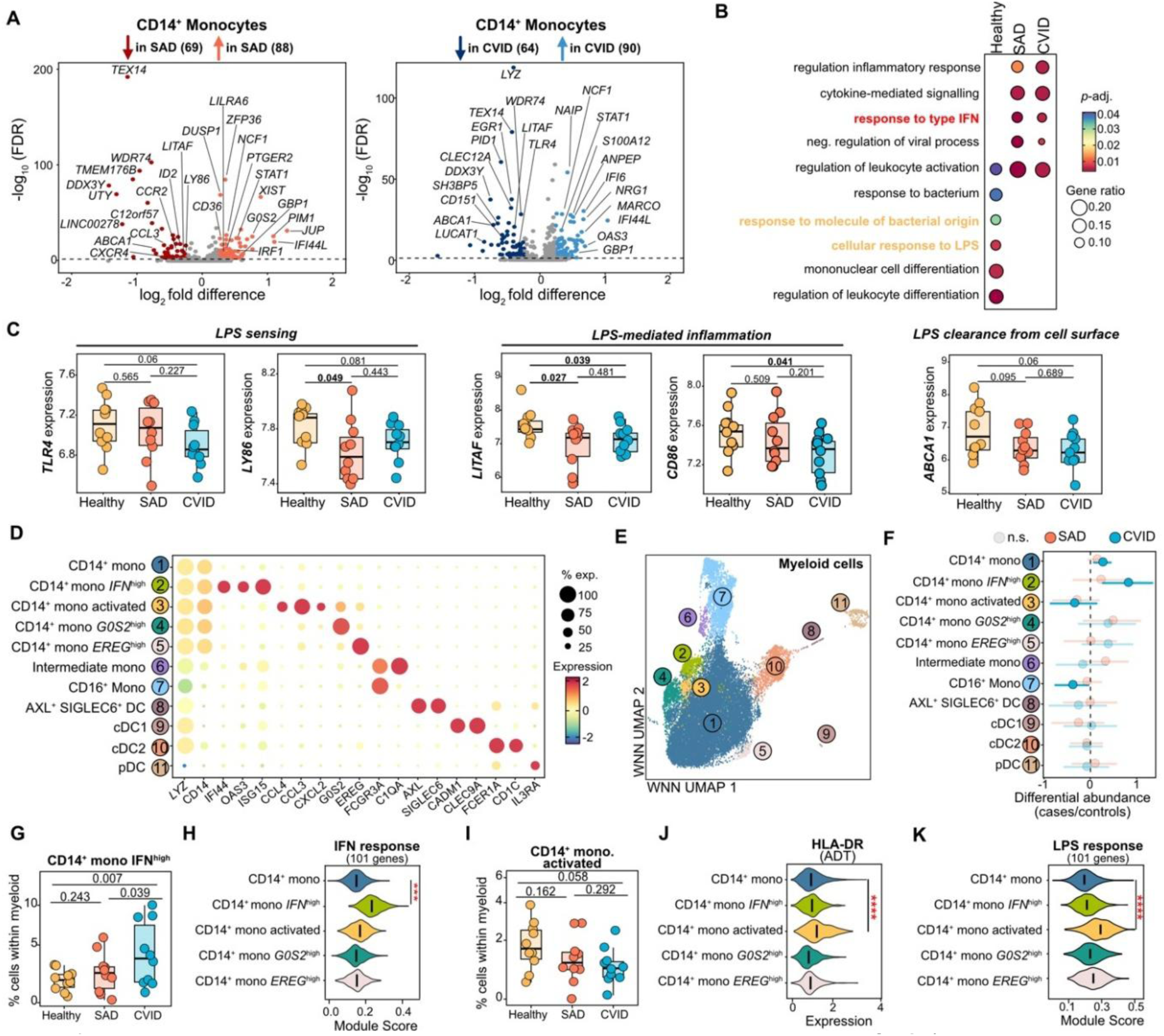
Altered lipopolysaccharide-associated gene expression and reduced CD14+ monocyte activation in SAD. **A)** Differential gene expression between cohorts in CD14+ monocytes. **B)** Gene ontologies for DEGs for CD14+ monocytes. **C)** Mean expression for genes associated with LPS responses. *p* denotes results of Student’s t-test. **D)** Marker gene expression for myeloid subsets. **E)** UMAP of 22,918 myeloid cell subsets. **F)** Relative cell type abundance between cases and controls. Error bars represent 95% credible interval; dashed black bars equal fold difference of 0.1. Opaque dots indicate significant results as per sccomp FDR-corrected *p*-value. **G)** Frequency of CD14+ IFNhigh monocytes. *p* values as per F. **H)** Expression of IFN-response gene module in myeloid subsets. *** denotes *p* < 1×10-200 (Wilcoxon signed-rank sum test). **I)** Frequency of activated CD14+ monocytes. *p* values as per F. **J)** Surface protein ADT expression of HLA-DR. **** denotes *p* < 2.22e-16 (Wilcoxon signed-rank sum test) **K)** Expression of LPS response module in CD14+ cells. *** denotes *p* < 2.02×10-68 (Wilcoxon signed-rank sum test).

### Diversity in T cell states as a feature of both health and immune dysfunction

We next examined CD4⁺ and CD8⁺ T cell states to understand if altered T cell responses may be relevant to understanding immune dysfunction in SAD. We identified 24 transcriptionally distinct populations, including five states of naïve CD4^+^ T cell (basal, IFN^high^, *SOCS2*^high^, *LOC105371574*^high^ and *LOC105376755*^high^); three states of CD4^+^ regulatory T cell (basal Treg, *FCRL3*^high^, *NR4A1*^high^); four populations of T helper cells (follicular helper T cells (Tfh), Th1, Th2, Th17); proliferating T cells; three states of naïve CD8^+^ T cell (basal, *LOC105371574*^high^, *S100B*^high^); three states of central memory CD8^+^ T cell (TCM; basal, *GATA*^high^ and *CCR9*^high^; three states of effector memory CD8^+^ T cell (TEMRA; basal, *PTGDS*^high^, inflammatory); CD4^-^CD8^-^double negative T (dnT) and cytotoxic CD4^+^ T cells (**Figure 3a**, **Figure S3a-c**).

**Figure 3.**
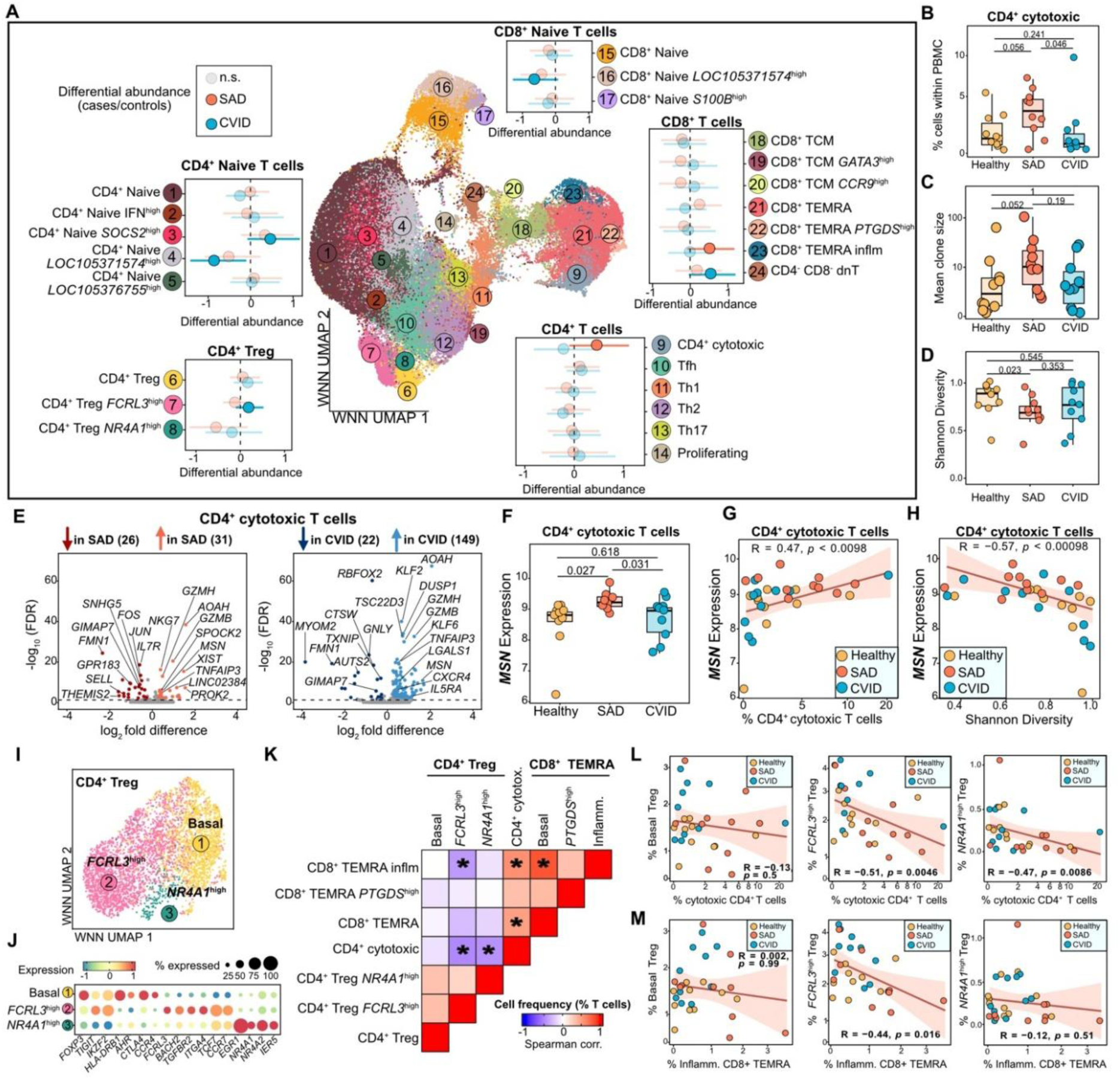
Expansion of cytotoxic CD4+ T cells and regulatory T cells defects in SAD. **A)** UMAP of 101,540 T cell states after high resolution clustering. Relative cell type abundance between cases and controls. Error bars represent 95% credible interval; dashed black bars equal fold difference of 0.1. Opaque dots and bars indicate significant results as per sccomp FDR-corrected *p*-value. **B)** Relative frequency of cytotoxic CD4+ T cells within PBMC. *p* values as per A. **C)** Mean clone size of cytotoxic CD4+ T cells. **D)** Normalised Shannon Diversity index for cytotoxic CD4+ T cells. **E)** Differential gene expression between cohorts for cytotoxic CD4+ T cells. **F)** *MSN* expression in CD4^+^ cytotoxic T cells (log2 CPM in cytotoxic CD4^+^ T cells). *p* denotes results of Student’s t-test. **G)** Correlation *MSN* expression (log2 CPM) in cytotoxic CD4^+^ T cells with cytotoxic CD4^+^ T cell abundance. **H)** As in G), for Shannon diversity index of cytotoxic CD4+ T cell clones. **I)** UMAP of 4,149 regulatory T cells states after high resolution clustering. **J)** Marker gene expression for regulatory T cell subsets. **K)** Correlation between the frequency of the T cell states by donor; * denotes *p*<0.05. **L)** Frequency of regulatory T cell subsets compared with cytotoxic CD4+ T cells, with Spearman correlation results shown. **M)** As in L) for inflammatory CD8+ TEMRA.

Some T cell states were highly variable between donors, including naïve CD4^+^ and CD8^+^ T cell states defined by expression of an uncharacterised long non-coding RNA gene *LOC105371574* that were more commonly observed in healthy donors, a *SOCS2*^high^ CD4^+^ naïve T cell state that was enriched in a small number of CVID patients, and a *PTGDS*^high^ CD8^+^ TEMRA population found in one CVID donor (**Figure 3a**, **Figure S3d-i**, **Table S3**). This highlights substantial inter-individual variability in T cell state composition, although the functional relevance in the context of disease is unclear.

### A unique expansion of cytotoxic CD4^+^ T cells in SAD

Cytotoxic CD4^+^ T cells are typically rare in healthy conditions but undergo clonal expansion during chronic infections and cancer^39^. Unexpectedly, we observed a unique enrichment of cytotoxic CD4^+^ T cells in SAD, but not in CVID (**Figure 3a-b**). To determine whether this expansion might reflect antigen-driven processes, we assessed clonal diversity within cytotoxic CD4⁺ T cells across cohorts and found that cytotoxic CD4^+^ T cells in SAD showed higher rates of clonal expansion (**Figure 3c-d**). Although individuals with SAD also had elevated frequencies of inflammatory CD8^+^ TEMRA cells, there was no difference in their rates of clonal expansion between cohorts (**Figure 3a, Figure S3j-k**) raising the possibility that there may be specific molecular features that mediate CD4⁺ but not CD8^+^ cytotoxic T cell expansion. Cytotoxic CD4^+^ T cells in SAD and CVID exhibited higher expression of genes involves with cytotoxicity (*GZMH*, *GZMB*, *NKG7*), LPS metabolism (*AOAH*), and T cell activation (**Figure 3e, Figure S3l**). CD4^+^ T cells in SAD had particularly high expression of *MSN* (encoding moesin; membrane-organizing extension spike protein) (**Figure 3e-f**) for which nonsense mediated mutations are associated with an X-linked primary immunodeficiency with T-cell lymphopenia^40^. Interestingly, higher *MSN* expression correlated with greater rates of clonal expansion of cytotoxic CD4^+^ T cells (**Figure 3g-h**), but not other effector or memory T cell states (**Figure S3m**).

Tregs regulate immune homeostasis by suppressing cytotoxic T cell responses and Treg-related defects have previously been reported in CVID^43^. We therefore considered if elevated cytotoxic CD4^+^ and inflammatory CD8^+^ TEMRA states in SAD could be related to altered frequencies or function of Tregs. We defined three Treg subsets, including *FCRL3*^high^ and *NR4A1*^high^ Treg states that were enriched for differential expression of genes associated with immune effector function, differentiation, and regulation of T cell activation (**Figure 3i-j**, **Figure S3n-o**). Intriguingly, individuals with fewer of these Treg subsets had increased cytotoxic CD4^+^ T cells and inflammatory CD8^+^ TEMRA (**Figure 3k-m**), suggesting alteration of Treg subsets may contribute to higher frequencies of inflammatory cytotoxic T cell states in SAD.

### Age-associated accumulation of NK cells and memory NK cell exhaustion in PAD

NK cells are critical mediators of early antiviral and antibacterial immunity, and reduced NK cell numbers have previously been associated with recurrent infections and other disease manifestations in CVID^22^. We observed increased frequencies of mature CD56^low^ NK cells in both CVID and SAD (**Figure 4a**). While NK cell frequency declined with age in healthy donors (with minimal other effects of age on immune cell frequencies), both CD56^high^ and CD56^low^ NK subsets increased with age in SAD and CVID (**Figure 4b**, **Figure S4a**). This suggests an age-related accumulation of mature NK cells in PAD, potentially as a result of recurrent infections^44^. We also observed that CD56^low^ NK cells in SAD and CVID expressed inflammatory and cytotoxic genes (e.g., *KLRC3*, *GZMA, CCL5, SLAMF7*) at higher levels (**Figure 4c**) with an enrichment for gene ontologies linked with cell killing and response to viruses (**Figure 4d**).

**Figure 4.**
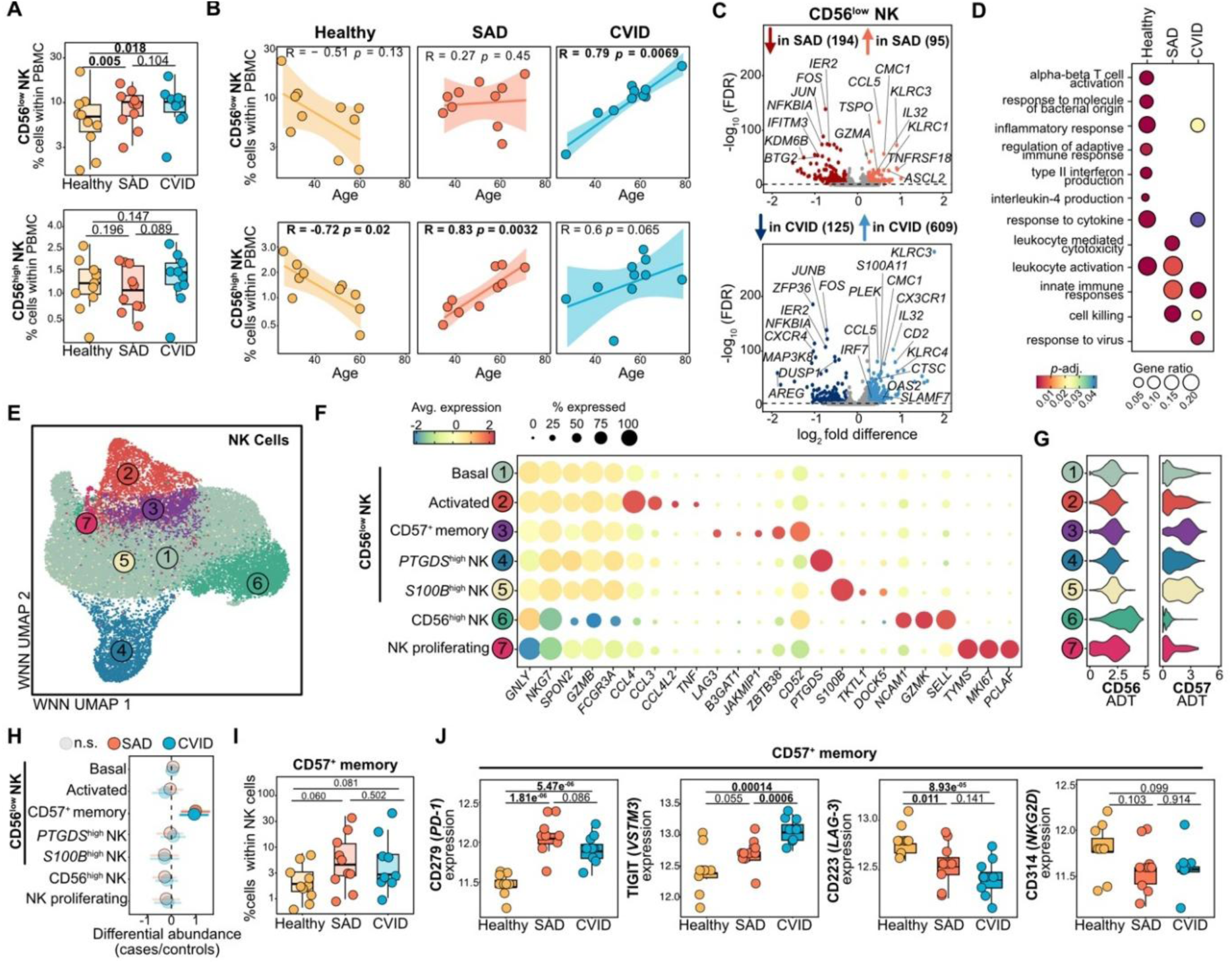
Age-associated accumulation and functional skewing of NK cell subsets in CVID and SAD. **A)** NK cell frequency relative to total PBMCs. *p* denotes FDR-corrected probability of compositional effect from sccomp. **B)** Association between NK cell frequency and age. Spearman’s correlation results shown. **C)** Differential gene expression for NK CD56low cells. **D)** Enriched gene ontologies for DEGs for CD56low NK cells (Healthy 214 genes, SAD 86 genes, CVID 565 genes). **E)** WNN UMAP of 17,518 NK cells. **F)** Marker gene expression for NK cells where dot size depicts frequency of cells a gene is detected in. **G)** Cell surface protein expression (ADT) for CD56 and CD57. **H)** Relative cell type abundance between cases and controls. Error bars represent 95% credible interval; dashed black bars equal fold difference of 0.1. Opaque dots and bars indicate significant *continued on next page* results as per sccomp FDR-corrected *p*-value. **I)** CD57+ memory NK cells relative to total NK cells. *p* values as per H. **J)** Expression of cell exhaustion genes in CD57^+^ memory NK cells (log2 CPM). *p* denotes Student’s t-test result.

We further identified seven distinct NK cell states, including five populations of CD56^low^ NK (basal, activated, CD57^+^ memory, *PTGDS*^high^ and *S100B*^high^), CD56^high^ NK and proliferating NK cells (**Figure 4e-g**). We found no major differences in NK cell subset frequencies between healthy controls and patients, except for an enrichment of CD56^low^ CD57^+^ memory NK cells in both SAD and CVID (**Figure 4h-i, Figure S4b, Table S3**). Higher expression of cell surface markers associated with cell exhaustion were also observed in CD57^+^ memory NK cells in SAD and CVID (**Figure 4j**), suggesting that prolonged inflammation or recurrent infections may increase exhaustion of NK memory. However, and somewhat surprisingly, we found that while CD57^+^ memory NK cells accumulated with age in CVID, they decreased in frequency in SAD (**Figure S4c-e**), raising the possibility of unique molecular or cellular defects in NK memory in SAD.

### Germinal centre-derived class-switched memory B cell subsets are reduced in PADs

In line with the importance of B cell-mediated antibody responses to infection and vaccination, there have been many examples of molecular and cellular dysfunction of B cell maturation in PADs^45,46^. This includes genetic mutations in key immune signalling transcription factors such as *NFKB1* and *NKFB2* that disrupt B cell activation in CVID^47,48^. However, little is known about the nature of B cell dysfunction in SAD^25^. While we did not observe as major defects of B cell maturation in SAD compared to CVID, with no significant difference in the proportion of CD27^+^IgD^-^ switched memory B cells when compared with healthy controls (**Figure 5a-b, Figure S5a**), there were widespread differences in gene expression within both naïve and memory B cells (**Figure S5b-c**). This included reduced expression of transcription factors involved in B cell development and maturation (*JUN*, *FOS*, *KLF2*, *BACH2*, *FOXO1*), immunoregulatory cell surface molecules involved with cell migration in lymphoid tissues (*CCR7*, *CD200*, *IL4R, P2RY8*, *S1PR4*, *TNFRSF13C*) and responses to stimulation (*FCMR*, *FCRL5*, *TLR9*) (**Figure 5c, Figure S5d**). Somewhat surprisingly, naïve and memory B cells in SAD exhibited higher expression of activation markers (*CD69*, *CD83*), interleukin-6 (*IL6*) and negative regulators of NFκB-dependent signalling (*NFKBIA*, *NFKBID*, *NFKBIZ*, *TNFAIP3*) (**Figure 5c, Figure S5c**). While some of these molecular differences were shared with CVID, some appeared to be specific to SAD and highlight potentially underappreciated molecular defects in B cell activation and/or maturation in SAD.

**Figure 5.**
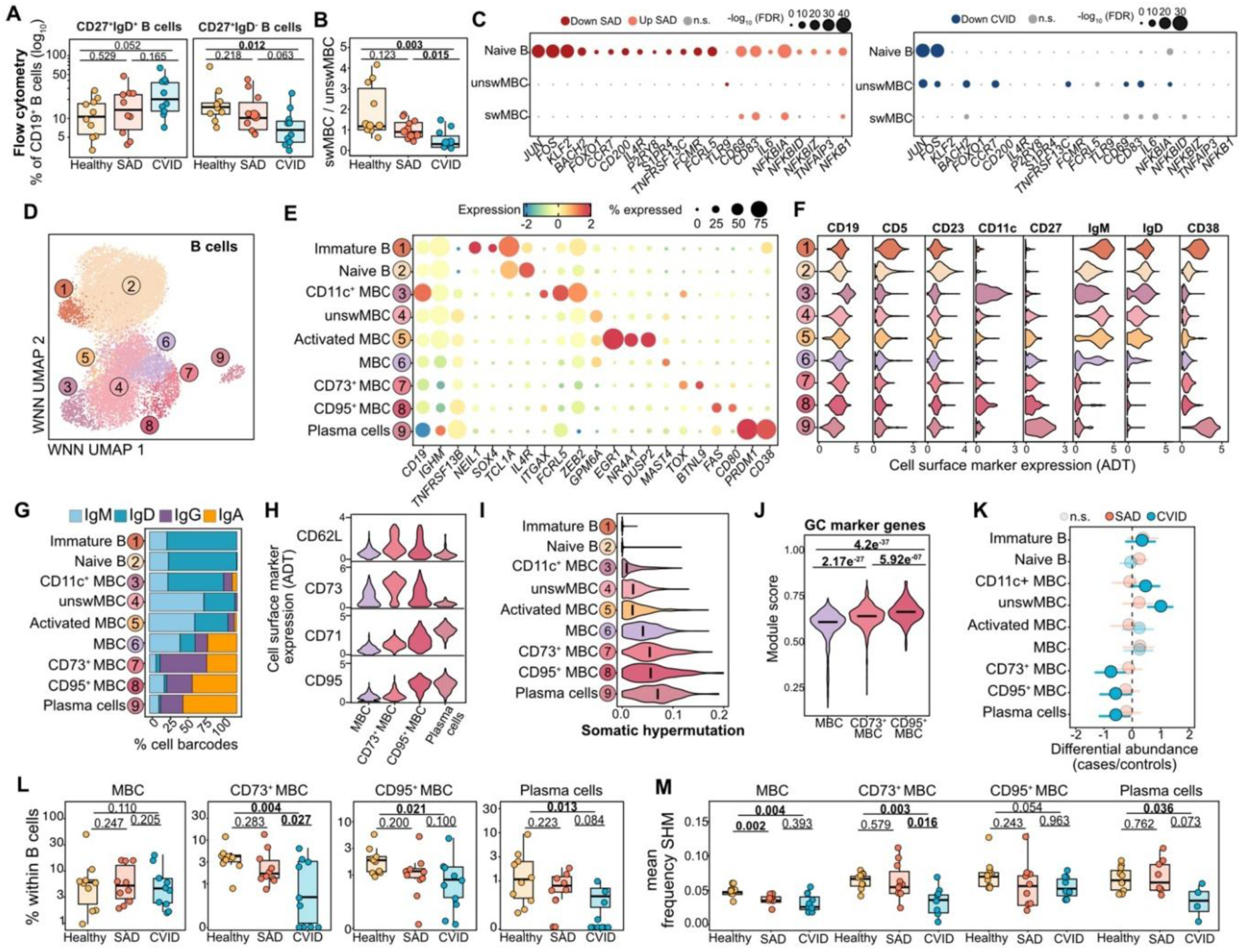
Dysfunctional B cell maturation in SAD and CVID. **A)** Proportion of CD27+ IgD+ B cells (unswMBC, left) and CD27+ IgD-B cells (swMBC, right) within CD19+ B cells across donors as determined by FACS. **B)** Ratio of proportions of swMBC and unswMBC across donors. **C)** Subset of DEGs identified across B cells between SAD and controls (left) or CVID and controls (right). Dot size depicts the -log10 FDR for a gene to be differentially expressed. **D)** UMAP of 14,392 B cells collected at baseline across donors and colored by their assigned high-level annotation. **E)** Mean expression top marker genes for B cell subsets. Dot size depicts frequency of cells a gene is detected in. **F)** Expression of selected cell surface markers across all B cell subsets. **G)** Frequency of antibody class within each B cell subset. **H)** Expression of selected cell surface markers within MBCs and plasma cells. **I)** Single-cell mean frequency of somatic hypermutation. **J)** Single-cell score of 250 GC marker genes derived from*49.* **K)** Estimated difference in cell type composition between cases and controls across B cell subsets. Error bars represent 95% credible interval; dashed black bars represent the minimum fold difference threshold of 0.1. Opaque dots and bars indicate significant results as per sccomp FDR-corrected *p*-value. **L)** Relative frequency of MBC, CD73+ and CD95+ MBCs and plasma cells relative to B cells. *p* values as per K. **M**) Mean somatic hypermutation frequencies for MBC, CD73+ and CD95+ MBCs and plasma cells. *p* values denote result of Wilcoxon signed-rank sum test.

To further explore this, we sub-clustered all B cells to resolve additional diversity of B cell states within SAD and CVID. Using a combination of unique gene expression, cell surface protein levels and antibody repertoires, we annotated nine distinct B cell states, including immature CD5^+^ B cells, naïve B cells, plasma cells, and six memory B cell states (**Figure 5d-g**). Within memory B cells, we identified three unswitched IgM^+^ memory populations (CD11c^+^ atypical, unswitched, activated). While CD11c^+^ atypical B cells have been described as age-related B cells^50^, we observed no correlation with age (**Figure S5e-f**). Instead, we found that individuals with elevated CD11c^+^ B cells also had higher frequencies of other unswitched memory B cell states (**Figure S5g-h**), potentially reflecting that CD11c^+^ B cells arise through similar maturation defects that bias towards unswitched memory in PAD. We also identified three predominantly class-switched IgM^-^IgD^-^ clusters that could be separated based on CD62L (*SELL*), CD73 (*NT5E*), CD71 (*TFRC*) and CD95 (*FAS*) expression (**Figure 5g-h**). CD73^+^ and CD95^+^ memory B cells exhibited comparable somatic hypermutation levels with plasma cells (**Figure 5i**) and genes enriched in these cells were elevated in tonsillar GC B cells^49^ (**Figure 5j, Figure S5i-k**). This is consistent with circulating CD73^+^ and CD95^+^ memory B cells likely having arisen through GC-dependent affinity maturation and could suggest they have more recently exited from lymphoid tissues. In keeping with known GC-dependent maturation defects in PAD, we found both SAD and CVID tended to have reduced frequencies of CD73^+^ and CD95^+^ memory B cells, and plasma cells (**Figure 5k-l, Figure S5l, Table S3**). Memory B cells and plasma cells in CVID had more marked reductions in somatic hypermutation rates than in SAD (**Figure 5m**, **Figure S5m**). This suggests that although participation in GC-dependent maturation may be less common in SAD, some B cells still achieve high affinity antibodies, whereas in CVID B cells are both less likely and less capable of developing high affinity antigen-specific responses.

### A spectrum of responses to Vi-polysaccharide vaccination in health and PAD

Our single-cell multi-omic atlas of SAD and CVID revealed unexpected differences across diverse immune cell subsets, which may contribute to their impaired ability to generate adequate antibody responses to polysaccharide antigens. To investigate this further, we vaccinated study participants with a typhoid Vi-polysaccharide vaccine (Typhim Vi) and monitored anti-Vi cellular and humoral responses (**Figure 6a**). While anti-Vi IgG titres in healthy controls increased upon vaccination (**Figure S6a-b**), we found that Australian-derived immunoglobulin replacement therapy products (of which 6/10 SAD and 6/10 CVID patients were receiving) contained moderate amounts of anti-Vi IgG (**Figure S6c-d**), which prevented accurate assessment of vaccine-induced humoral responses in the CVID and SAD cohorts. However, analysis of Vi-specific antibody-secreting cells at day 7 post vaccination (previously shown to be the peak of antibody-secreting cell responses^51^) for healthy donors showed rapid antigen-specific responses, while there were more limited responses in SAD and minimal responses in CVID (**Figure 6b**). Overall, this revealed a dynamic range of early responses to vaccination spanning health and PAD (**Figure 6c**).

**Figure 6.**
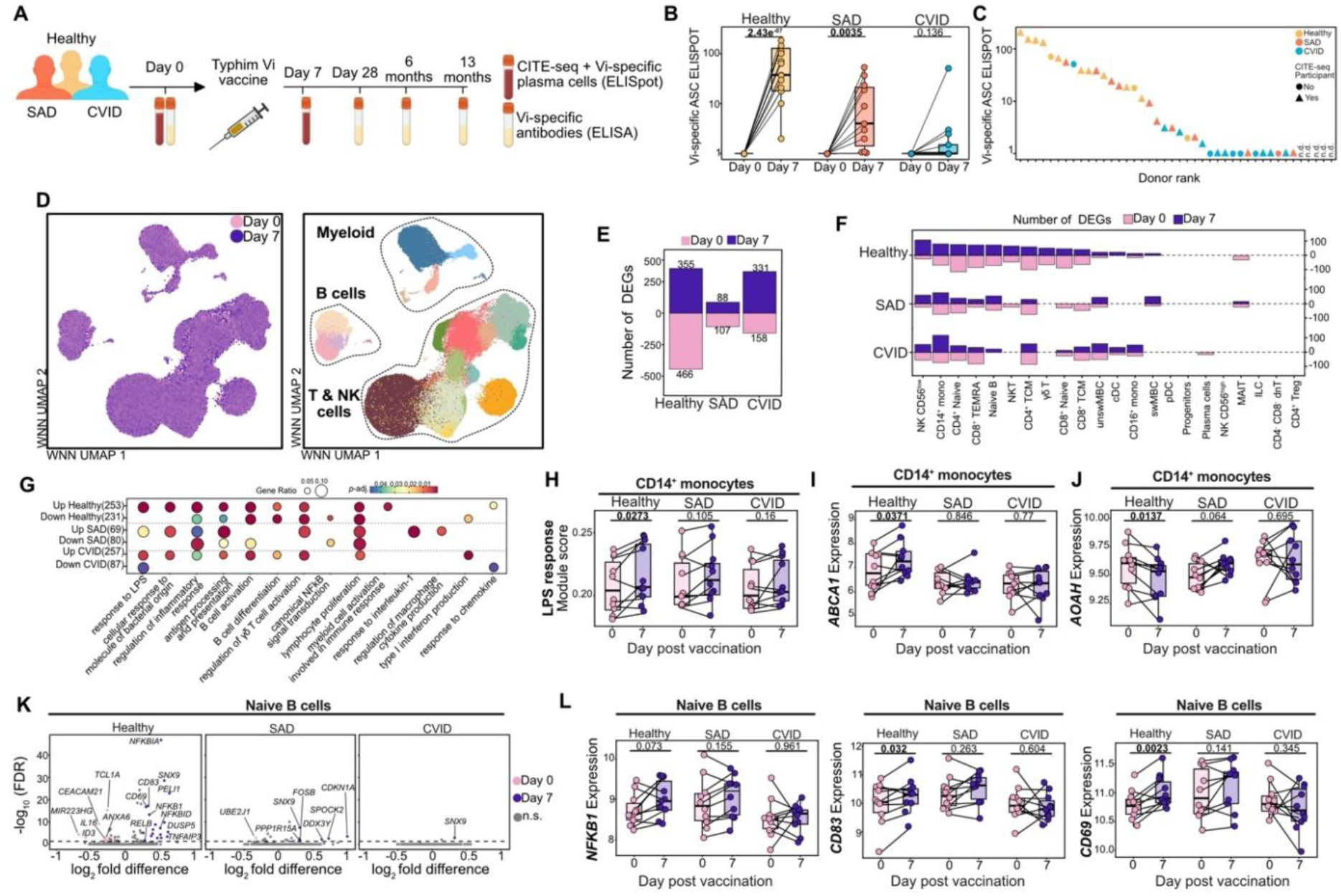
Innate immune sensing and B cell activation defects in PAD following vaccination. **A)** Vaccination study design. After vaccination with a Typhim Vi polysaccharide vaccine, samples were collected at different timepoints. **B)** Frequency of antigen-specific antibody secreting cells (ASCs) via ELISpot at baseline and 7 days post-vaccination. *P* values denote result of Student’s t-test. **D)** UMAP of 322,174 single-cells coloured by day cell type annotation. **E)** Number of DEGs between vaccination timepoints. **F)** Number of DEGs between vaccination timepoints across cohorts for cell types. **G)** Enriched gene ontologies for DEGs d in F), with number of DEGs reported within parentheses. **H)** Mean expression of 101 genes associated with responses to LPS before and after vaccination within CD14+ monocytes. *p* values denote result of paired Wilcoxon signed-rank sum test. **I)** Mean expression of *ABCA1* within CD14+ monocytes between vaccination timepoints. *p* values denote result of paired Student’s t-test. **J)** Mean expression of *AOAH* within CD14+ monocytes between vaccination timepoints. *p* values denote result of paired Student’s t-test. **K)** DEGs identified in naïve B cells between baseline and 7 days post-vaccination for each cohort. **L)** Mean expression of *NFKB1*, *CD83* and *CD69* in naïve B cells between timepoints. *p* values denote result of paired Student’s t-test.

### Dynamic monocyte and B cell transcriptional responses to vaccination absent in PAD

To resolve the dynamics of early immune responses to vaccination with the Vi polysaccharide antigen, we performed CITE-seq on PBMCs isolated after vaccination at day 7 and compared them with our previous analysis of PBMCs collected before vaccination (day 0) (**Figure 6d**). We found widespread differences in gene expression of circulating immune cells following vaccination in healthy donors (**Figure 6e**), with many diverse cell types and subsets responding to vaccine challenge (**Figure 6f-g**). While in some cases we observed PAD-specific responses, such as variably elevated IFN-related gene expression in CD14^+^ monocytes of CVID patients (**Figure S6e-f**), most gene expression changes following vaccination were only apparent in healthy donors. For example, CD14^+^ monocytes in the healthy control cohort consistently up-regulated genes associated with molecular responses to LPS while this was not observed in SAD or CVID (**Figure 6h**), such as the macrophage ATP-binding cassette transporter A1 (*ABCA1*) that is required for the removal of cell-associated LPS and ensuring macrophage responsiveness^41^ (**Figure 6i**).

In contrast to healthy controls, we found that SAD patients up-regulated *AOAH* expression after vaccination with the Vi polysaccharide (**Figure 6j**). Given a role for *AOAH* in de-acylation and inactivation of LPS^31^, which can be present in Vi-polysaccharide vaccine preparations and contribute to their immunogenicity^52^, this differential expression in SAD could contribute to attenuated *TLR4* signalling and dampen downstream inflammatory responses that are a typical feature of immune responses to this vaccine. Finally, we found rapid transcriptional activation in circulating naïve B cells after vaccination only in healthy donors, with increased expression of genes involved in B cell activation and NF-kB-dependent signalling (*NFKB1, RELB, TNFAIP3, DUSP5, CD83, CD69*) (**Figure 6k-l, Figure S6g**). These differences in early post-vaccination gene expression suggest that reduced monocyte-mediated immune stimulation may impair B cell activation and contribute to diminished antigen-specific responses in PAD.

### Circulating proliferative monocyte and CD95^+^ memory B cell states as an early correlate of Vi-polysaccharide vaccine response

Although adaptive immune responses to vaccination are predominantly driven by processes restricted to lymphoid tissues such as GC-dependent maturation of antigen-specific B cells, previous studies have shown that some of these processes can be captured by transient changes in immune cell abundances in peripheral blood, thus providing an accessible readout of cellular responses to vaccination in PAD. We therefore examined whether there were any changes in the relative abundance of 58 immune cell states after vaccination (**Figure 7a**).

**Figure 7.**
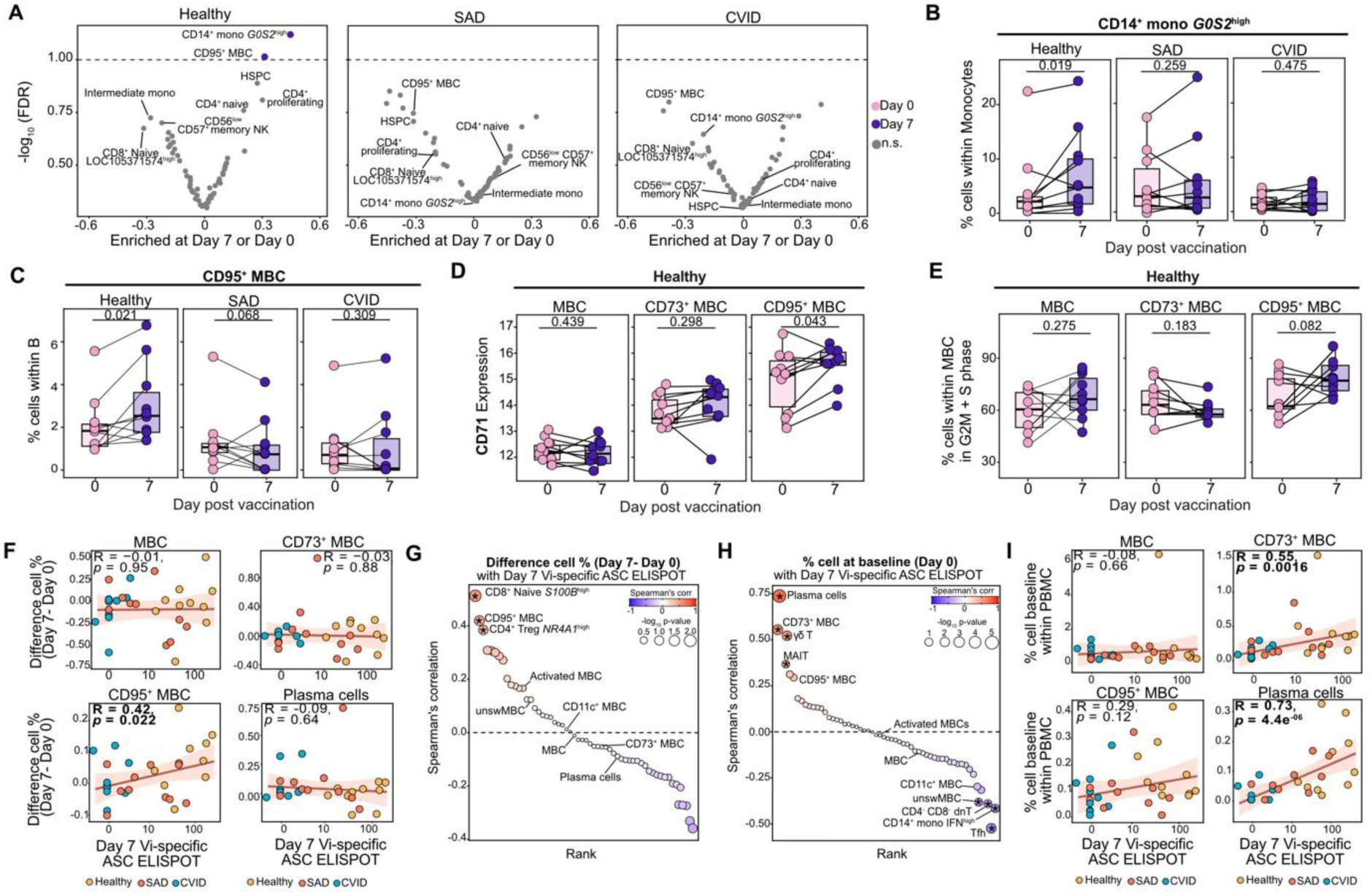
Rapid expansion of circulating monocytes and CD95+ activated memory B cells after Vi-polysaccharide vaccination. **A)** Differential cell type abundance between vaccination timepoints. **B)** Frequency of *G0S2*high CD14+ monocytes within monocytes between timepoints. *p* denotes FDR-corrected probability of effect between vaccination timepoints by sccomp. **C)** Frequency of CD95+ memory B cells (MBCs) within B cells between timepoints. *p* as in B. **D)** Expression of cell surface marker CD71 (log2) across MBCs in controls. *p* values denotes paired Student’s t-test. **E)** Percentage of G2M/S phase cells within controls between timepoints. *p* as in D. **F)** Difference in cell frequency between timepoints compared with Day 7 Vi-specific ASCs. **G)** Spearman’s correlation of cell type frequency changes (Day7 - Day0) and Vi-specific ASCs at day 7, where dot size reflects significance of correlation. * denote *p*-value <0.05. **H)** Spearman’s correlation of cell type frequency at baseline (Day 0) and Vi-specific ASCs at day 7, where dot size reflects significance of correlation. * denote *p*-value <0.05. **I)** Relative MBC and plasma cell frequency at day 0 compared with Vi-specific ASCs at day 7. Spearman’s correlation results are shown.

While we had previously found differences in Vi-specific antibody-secreting cell responses following vaccination, we did not observe any significant changes in total plasmablasts or plasma cell frequencies (**Figure S7a**). Instead, on day 7 post-vaccination, we identified increased frequencies for a CD14^+^ monocyte state (that we annotated as *G0S2*^high^ CD14^+^ monocytes) and CD95^+^ memory B cells (**Figure 7a-c**, **Figure S7b-c**). Notably, both cellular responses were reduced, absent or more variable in SAD and CVID (**Figure 7a-c**), suggesting that they could be related to the formation of Vi-specific responses. We first examined the *G0S2*^high^ CD14^+^ monocytes, which compared with other CD14^+^ monocyte states had high expression of genes involved in myeloid cell activation, inflammation and monocyte differentiation (**Figure S2c-d**). In addition to inflammatory cytokines such as *IL1B* (IL-1β) and *CXCL8* (IL-8) that can be up-regulated in response to TLR-dependent signalling, these cells also expressed higher levels of pro-survival *BCL2A1* and G0/G1 switch gene 2 (*G0S2*) that was first described in peripheral blood mononuclear cells transitioning into cell cycle and DNA replication^53^(**Figure S2c**). However, we did not find evidence for increased proliferative signatures post-vaccination (**Figure S7d-e**). Vaccine-dependent increases in this activated monocyte state were unrelated to the magnitude of the antigen-specific response (**Figure S7f**), suggesting that this cellular response might be more relevant to inherent molecular defects in endotoxin- or polysaccharide-based innate immune sensing between the cohorts than as a key mediator of antigen-specific immune responses.

We next investigated the expansion of CD95^+^ memory B cells at day 7 post-vaccination, predominantly in healthy participants (**Figure 7c**, **Figure S7c**). We previously observed that this class-switched and high-affinity memory B cell population had high expression of GC-associated genes, potentially suggesting recent egress from lymphoid tissues. CD95^+^ memory B cells had high expression of CD71 (*TFRC*) which functions as a major iron transporter required for rapid cellular expansion and is associated with activated and proliferating B cells in the circulation following infection or vaccination^54^ (**Figure 7d**). Indeed, we observed a higher frequency of memory B cells transitioning into S and G2/M phases of the cell cycle at day 7 post-vaccination (**Figure 7e**), consistent with increased proliferation of these cells. Excitingly, the vaccine-dependent increases in CD95^+^ memory B cells frequencies (but not other memory populations) positively correlated with the magnitude of antigen-specific vaccine responses across the spectrum of health and immunodeficiency in our cohort (**Figure 7f-g**). When we examined whether changes in any of the other 57 immune cell states in our vaccination atlas correlated with the magnitude of the day 7 antigen-specific response, we also identified significant increases in a naïve *S100B*^high^ CD8^+^ T cell state and activated *NR4A1*^high^ CD4^+^ Treg cells (**Figure 7g, Figure S7g-h**), which likely reflect bystander activation. Finally, we examined whether frequencies of these and other cell types measured prior to vaccination could be used to predict the magnitude of the antigen-specific response to the Vi polysaccharide vaccine (**Figure 7h**). Across the three cohorts we observed that high frequencies of plasma cells, CD73^+^ memory B cells and non-conventional T cells prior to vaccination were associated with greater Vi-specific antibody-secreting cell responses (**Figure 7h-i, Figure S7i**).

## DISCUSSION

The molecular and cellular mechanisms underlying inter-individual variability in responses to polysaccharide vaccines remain incompletely understood, despite their clinical use in diagnosing PADs, such as SAD. Using longitudinal single-cell multi-omics, we show that impaired vaccine responsiveness is associated with coordinated defects spanning both innate and adaptive immunity. We identify dysregulated monocyte activation linked to LPS sensing, accumulation of inflammatory and exhausted lymphocyte states, and reduced frequencies of GC-associated memory B cells across PADs. Although SAD has historically been difficult to classify, our data place it within a broader spectrum of systemic immune dysfunction with a unique expansion of clonally derived cytotoxic CD4⁺ T cells. Early Vi-polysaccharide vaccine responses were associated with dynamic activation of monocytes and B cells together with expansion of proliferative CD95⁺ memory B cells, responses that were attenuated or absent in PAD. Together, these findings provide a cellular framework for understanding defective polysaccharide vaccine responses and redefine SAD as a broader disorder of immune dysregulation rather than an isolated defect in antibody production.

This is the first study to provide an in-depth characterization of immune cell types and molecular signatures of SAD, a poorly understood yet commonly diagnosed PAD. Without genetic or gold-standard diagnoses, SAD patients represent a heterogenous population for which only limited studies have directly compared them with more well studied conditions such as CVID^27^. While many cellular defects were shared between SAD and CVID, including reduced non-conventional T cells, activated CD14^+^ monocytes, plasma cells and high-affinity memory B cells, these cellular differences were generally less pronounced and more variable in SAD. While it is possible that some SAD individuals represent evolving CVID^55^, we identified examples of immune cell dysfunction that were unique to SAD, even with the modest size of our cohort. Larger future studies of SAD patients in comparison to other PADs and healthy cohorts will be beneficial to further quantify and resolve the immunological and molecular features that underlie their reduced ability to mount long-lived vaccine responses to polysaccharides.

The expansion of cytotoxic and memory-associated lymphocyte populations, including clonally-expanded cytotoxic CD4⁺ T cells and inflammatory CD8⁺ TEMRA cells was a striking feature of SAD. CD8⁺ TEMRA cells are a well-established terminal effector memory population associated with chronic antigen exposure, viral infection, and immune exhaustion, with relatively defined developmental trajectories and effector functions. In contrast, cytotoxic CD4⁺ T cells remain more heterogeneous and less clearly understood^56^. Although increasingly recognised as important mediators of antiviral immunity and immune surveillance, cytotoxic CD4⁺ T cells have also been associated with tissue damage and disease severity in autoimmune and inflammatory disease^57,58^. It therefore remains unclear whether their expansion reflects a protective adaptive response or a marker of chronic immune dysregulation and inflammation in SAD. While cytomegalovirus (CMV) infection or re-activation has been implicated in the expansion of cytotoxic CD4⁺ T cells^59^, the lack of expanded cytotoxic CD4^+^ T cells in CVID where chronic CMV reactivation has been well established^29,60^ suggests other mechanisms are involved. We also observed a negative correlation between cytotoxic CD4^+^ T cell abundance and *NR4A1*^high^ or *FCRL3*^high^ Treg cell states. NR4A receptors are essential for Treg differentiation and maintenance of T cell homeostasis^61^. *Nr4a1*^-/-^ mice exhibit exaggerated cytotoxic T cell responses in response to bacterial infection^62^ and higher FCRL3 expression has been associated with reduced Treg activation and regulatory potential^63,64^. It is also possible that T cell-intrinsic mechanisms could explain the expansion of cytotoxic T cells in SAD. For example, higher moesin (*MSN*) expression in SAD correlated with higher rates of cytotoxic CD4⁺ clonal expansion, consistent with the role of moesin in T cell activation^65^. Of note, mutations in *MSN* lead to an X-linked recessive genetic disease whereby patients present with hypogammaglobulinemia, poor responses to polysaccharide antigens and lymphocytopenia^41^. While our whole-genome sequencing of the SAD cohort did not identify any strong candidates for a genetic basis of this type of molecular defect, this will be an important area for future research.

SAD is clinically defined by impaired antibody responses to polysaccharide-based vaccines, and our single-cell resource provided an opportunity to identify molecular pathways and cell states that may mediate this. Our data support dysregulation of innate immune metabolism, sensing, and response to LPS pathways as features of impaired vaccine responses for SAD and CVID. At steady-state, altered expression for genes associated with cellular responses to LPS, including LPS sensing (*TLR4*, *LY86*) and LPS-mediated inflammatory responses (*CD86, LITAF*), were observed. We defined a rare activated CD14^+^ monocyte state which was less abundant in SAD and CVID that appeared to mediate these LPS-related functions. It is unclear if the difference between cohorts relates to a difference in the functional potential of these cells to become activated or unknown developmental mechanisms. While healthy individuals responded to vaccination with upregulation of LPS-responsive genes in CD14^+^ monocytes, this was not observed in SAD and CVID. LPS-specific responses have previously been reported following Vi-polysaccharide vaccination and is likely due to LPS contamination of the Vi-polysaccharide antigen^66,67^. As blood samples were collected at 7-days post-vaccination (to capture the peak of circulating Vi-specific antibody secreting cells in peripheral blood)^67^, we cannot exclude that monocytes at earlier or later post-vaccination timepoints in SAD and CVID individuals are still capable of this response. However, we propose that an inability to sense or metabolise LPS contaminants in PAD reduces the adjuvant-like effect of LPS which affects the immunogenicity of the Vi-polysaccharide vaccine. This could contribute to reduced antigen-specific responses. In support of this hypothesis, the elevated expression of acyloxyacyl hydrolase (*AOAH*) in SAD could increase de-acylation of LPS and reduce LPS-associated immune stimulation and downstream antibody production^31^.

While multiple studies have investigated vaccine-induced immune responses and memory in healthy individuals^1–3,68^, less is known about the molecular and cellular features after vaccination in CVID^69–71^ and SAD. As most SAD and CVID patients included in our longitudinal Vi-polysaccharide vaccine challenge received immunoglobulin replacement therapy products which contained moderate amounts of anti-Vi antibodies, we could not use anti-Vi IgG titres to classify vaccine responders and non-responders. We therefore used Vi-specific antibody secreting cells detected 7-days post-vaccination, as our measure of antigen-specific vaccine response. While in some settings antibody secreting cell responses do not correlate with peak antibody titres, longer-term serum antibodies and Vi-specific antibody secreting cells analysis at day 7 were reasonably well correlated. While CVID patients failed to mount Vi-specific responses, the responses in SAD were more variable. How this relates to clinical outcomes in SAD patients is not yet clear. We were, however, able to ascribe several differences in baseline (day 0) cell type frequencies as strong indicators of an individual’s ability to respond to future immune challenges. Most notably plasma cells and class-switched memory B cells positively correlated with vaccine-specific antibody secreting cell responses. Conversely, higher frequencies of unswitched memory B cells and inflammatory monocytes were negative predictors of vaccine response, the latter of which is consistent with studies showing abundant pro-inflammatory innate immune cell states are associated with weak antibody responses to a wide range of infections^2^.

Long-lived and high-affinity antibody responses to vaccination typically arise via GC-dependent affinity maturation in secondary lymphoid organs like the spleen, lymph nodes and tonsil and recent efforts have made significant progress in defining these post-vaccination processes with longitudinal fine-needle aspirate studies^72,73^. Although our study was limited to peripheral blood, we were able to identify dynamic and rapid responses in B cell activation, via NF-kB signalling pathway in naïve B cells, and expansion of a CD95^+^ memory B cell population at day 7 post-vaccination. These differences were absent or abrogated in SAD and CVID and correlated with the magnitude of antigen-specific responses. Class-switched memory B cells expressing CD95^+^ have been proposed to represent early emigrants of the GC^74^ primed for rapid differentiation into antibody-secreting cells^74,75^, and we provide further evidence that these cells share transcriptional and antibody repertoire properties consistent with recent exit from the GC. Increased frequencies of circulating CD95^+^ memory B cells have recently been reported in a large cohort of more than 300 healthy adults following seasonal influenza vaccination^3^ and in *Plasmodium falciparum* infections^75^. CD95^+^ memory B cells also express CD62L that was previously associated with antigen-experienced, GC-derived cells after SARS-CoV2 mRNA vaccination^76^, as well as CD71 which marks a transiently-activated population of memory B cells in the blood following viral infection or influenza vaccination^77^. Fittingly, we identified evidence of increased proliferation in CD95^+^ memory post-vaccination, albeit with transcriptional markers of cell division rather than phenotypic measures. In contrast to viral and mRNA-based vaccination strategies, polysaccharide antigens including the Vi-polysaccharide vaccine are thought to be largely T cell-independent and thus not rely upon GC-dependent processes, raising interesting questions about how CD95^+^ memory B cells are activated or generated in response to vaccine challenge. Although the Vi-polysaccharide vaccine is administered intramuscularly, vaccine-induced anti-Vi IgA responses and gut-homing IgA+ plasma cells have been observed post-vaccination (day 7), suggesting that a memory B cell population primed in mucosal lymphoid sites could occur in typhoid-naïve individuals^67,78,79^. Further studies involving sampling of secondary lymphoid tissues, rather than peripheral blood, would help to elucidate the exact origin of these CD95^+^ memory B cells following Vi-polysaccharide vaccination.

Together, our findings place SAD within a broader spectrum of immune dysfunction and identify cellular and molecular features of effective polysaccharide vaccine responses, including expansion of CD95⁺ memory B cells that may serve as a surrogate marker of vaccine responsiveness and provide a foundation for improved cellular diagnostics for PAD. More broadly, approaches based on cellular immune responses rather than antibody titres may enable more accurate assessment of humoral immunity in patients receiving immunoglobulin replacement therapy, where conventional serological measurements are difficult to interpret.

## ACKNOWLEDGEMENTS

The authors thank all volunteers for their participation and for providing samples essential to this study. The authors thank Daniela Zalcenstein, Casey Anttila, Rory Bowden, Peter Hickey and members of the WEHI Advanced Genomics Facility and gratefully acknowledge the WEHI Flow Cytometry Facility for their support and assistance in this work. Some figures were made with the assistance of BioRender.

## AUTHOR CONTRIBUTION

Conceptualization, H.W.K., C.J., D.V., M.H; data curation, H.W.K., D.V., C.J., A.J., K.D.; formal analysis, D.V., A.Q., J.L., J.C., C.Y.S., C.J., H.W.K.; funding acquisition, C.J., H.W.K., J.J.B., S.O., C.S.; investigation, D.V., H.W.K., C.J., J.L., A.Q., Z.K.T.; methodology, H.W.K., D.V., C.J., M.H., J.J.B., S.O., C.S., S.C., J.C., Z.K.T., J.L., A.Q.; project administration, C.J., H.W.K., D.V., A.J., K.D., S.T., S.S., S.C., J.J.B., J.C., C.S., S.O.; resources, D.V., C.J., J.L., A.Q., E.C.L., M.M.; supervision, H.W.K., C.J., J.J.B.; visualization, D.V.; writing – original draft, D.V., H.W.K., C.J.; writing – review & editing, D.V., A.Q., J.L., Z.K.T., A.J., S.S., S.T., K.D., E.C.L., M.M., M.H., S.C., J.J.B., J.C., S.O., C.S., C.J., H.W.K.

## METHODS

### Participants and Study Procedures

The study protocol was approved by the Sponsor (Royal Melbourne Hospital) and the Melbourne Health Human Ethics Committee (HREC/73845/MH-2021). The study was registered with the Therapeutics Good Administration (Clinical Trial Notification scheme), and the Australian New Zealand Clinical Trials Registry (ACTRN12621001059853). The study was conducted in accordance with the ethical principles of the Declaration of Helsinki.

Adult SAD (n=11), CVID (n=16) and healthy volunteers (n=15) were recruited through the Royal Melbourne Hospital and Alfred Health, two tertiary immunology referral centres in Victoria, Australia from January 2022 to May 2023. A subset of participants (n=10 per group) was included for single-cell analysis based on availability and quality of PBMC samples. CVID and SAD patients were receiving immunoglobulin replacement therapy for ≥6 months prior to enrolment, and as such fulfilled diagnostic criteria as per the Australian National Blood Authority Qualifying Criteria, which reference the European Society of Immunodeficiencies diagnostic criteria for CVID and the American Academy of Allergy, Asthma and Immunology 2015 Practice parameter for the diagnosis and management of primary immunodeficiency for SAD^80^. At the time of enrolment, two participants were receiving immunomodulatory therapies (one CVID patient received dupilumab [anti-IL-4Rα monoclonal], and one SAD patient receiving ustekinumab [anti-IL-12/IL-23 monoclonal]). Exclusion criteria for healthy volunteers included Vi-polysaccharide vaccination <3 years prior to enrolment; immunosuppressive treatment or active malignancy. General exclusion criteria for all groups included B cell depletion treatment within 2 years of enrolment and pregnant/breast-feeding women.

### Vi-polysaccharide vaccine challenge and anti-Vi IgG quantification

Written informed consent was obtained for all participants at the time of study enrolment. A single dose of Vi-polysaccharide vaccine (TYPHIM Vi™; Sanofi Pasteur, Lyon France), containing 25mcg of purified *Salmonella* Typhi capsular Vi-polysaccharide, was administered intramuscularly on day 0 (baseline). Follow up visits were performed at 7 days and 1, 6 and 13 months post-vaccination. Serum samples collected at baseline and 1, 6 and 13 months post-vaccination were stored at -80°C prior to analysis. Concentrations of anti-Vi IgG were measured directly from commercial immunoglobulin replacement therapy products; two different batches of Australian blood donor derived Intragam® 10 (CSL Behring, Broadmeadows, Australia), and international blood donor derived products Privigen® (CSL Behring, Broadmeadows, Australia) and Octagam® 10% (Octapharma, Pyrmont, Australia). Immunoglobulin replacement therapy products were pre-diluted to a concentration of ∼10g/L and were serially diluted to investigate for possible antibody excess effects. Anti-Vi IgG titres were measured using the VaccZyme™ Anti-*S.* Typhi Vi human IgG EIA (The Binding Site, Birmingham UK) as per the manufacturer’s instructions. Plates were read using a VersaMax microplate reader (Bio-Strategy, Tullamarine Australia) with Soft Max Pro. 6 software. Samples with undetectable antibody titres, less than the 7.4 U/ml lower limit of detection, were assigned values of 3.7 U/ml for the purposes of calculating fold change in titre.

### Vi-specific antibody secreting cell quantification by ELISpot

PBMCs were separated and cryopreserved from freshly isolated blood samples collected at baseline and 7-days post-vaccination. Antigen-specific ASCs were measured using an enzyme-linked immunosorbent spot (ELISpot) assay, as previously described^81^. Briefly, 96-well Multiscreen filter plates (Merck Millipore, Burlington, USA) were coated with 10ug/ml of Vi capsular polysaccharide (NIBSC 12/244, Lot 2039, NIBSC, Potters Bar, UK). Thawed PBMCs were resuspended in complete media (RPMI supplemented with 10% heat inactivated foetal bovine serum, 1% L-glutamine, 1% penicillin/streptomycin, 1% MEM Non-essential amino acids, 1% sodium pyruvate, 0.1% β-mercaptoethanol) and incubated at 37°C with 5% CO2 overnight on pre-coated 96-well plates at a concentration of 2.5×10^5^ PBMCs per well. Plates were developed with goat anti-human IgG alkaline phosphate conjugates (Calbiochem/Merck, Burlington, USA) and substrate development kits (Bio-Rad Laboratories Ltd, Watford, UK) the following day and counted using the AID *i*Spot, software (v7.0).

### Single-cell library preparation, sequencing and alignment

PBMC samples were stained with Hash Tag Oligonucleotides (HTOs) and a panel of 130 Antibody-Derived Tags (ADTs) using TotalSeq^TM^ antibodies (BioLegend). All 60 samples were then pooled and split into 5 batches (12 samples each). Each batch comprised two individuals from each cohort, with day 0 and day 7 samples labelled separately with unique HTOs (**Table S4**). For each batch, GEX, ADT and HTO and VDJ libraries were prepared at the Advanced Genomics Facility at the Walter and Eliza Hall Institute using the 10X Genomics Chromium single-cell 5′ reagent kit v3 according to manufacturer’s protocol. Libraries were then sequenced on a NovaSeq X Plus (PE150). Bcl2fastq (v2.20.0; Illumina) was used to convert and demultiplex CITE-seq and VDJ-seq BaseCall files into FASTQ files. Cellranger mkref and cellranger mkvdjref (v8.0.1; 10x Genomics) were used to build the hs1/T2T-CHM13v2.0 reference genome respectively for the CITE-seq and VDJ-seq data using the GCF_009914755.1_T2T-CHM13v2.0 annotations and hs1 genome build downloaded from UCSC. Cellranger multi and cellranger vdj (v8.0.1; 10x Genomics) were respectively used to align the CITE-seq and VDJ-seq libraries to the hs1 reference genome and to generate the cell barcode-gene UMI count matrices.

### CITE-seq demultiplexing, quality control, batch integration and cell clustering

Gene expression count matrices for each sequencing batch were separately processed with Seurat (v.5.4.0)^82^ in R (v. 4.4.1). Briefly, HTODemux^83^ (assay = “HTO”, positive.quantile = 0.99, kfunc=’clara’) was used to assign the donor and vaccination timepoint for each single cell based on cell hashtags and to remove both doublets and negative droplets. ScDblFinder^84^ with default parameters was then used to further identify and remove heterotypic doublets. Resulting singlets were then filtered to retain high-quality cells based on the number of genes per cell (between 200 and 9000), number of UMIs per cell (between 500 and 7000) percentage of mitochondrial reads per cell (0 to 5%), and whether the cell was predicted as an outlier by the isOutlier function of the scater R package^85^ based on the number of ADTs. Preprocessed Seurat objects for each of the 5 sequencing batches were then merged. Gene expression (RNA) data was then normalised using SCTransform^86^, regressing out the percentage of mitochondrial and ribosomal genes and specifying 4,000 variable features. The ADT data was normalised using the centered log ratio normalization before scaling and principal component analysis (PCA) reduction. Batch correction was performed separately on ADT and RNA using the RPCA integration in Seurat. Principle component analysis for ADT and RNA assays were run separately prior to using the weighted nearest neighbors (WNN) approach^87^ for uniform manifold approximation and projection (UMAP) dimensionality reduction and clustering (50 and 25 PCs for RNA and ADT respectively).

### Cell type clustering and annotation

FindClusters with default parameters and resolution 0.4 was used for cluster identification and then FindAllMarkers with default parameters was used to manually annotate clusters based on the expression of marker genes and ADTs. High-resolution cell clustering and annotation was achieved by subsetting each main cell lineage (i.e., B, T, NK and myeloid cells) and then performing lineage-specific batch integration via RPCA, WNN clustering and UMAP reduction. Marker genes for clusters were identified using FindMarkers or FindAllMarkers, and a combination of known marker genes for immune cell subsets and *de novo* marker gene identification was used to assign high resolution cluster cell type/state annotations.

### Single-cell BCR and TCR repertoire analysis

Cellranger single-cell VDJ datasets for BCR and TCR from all_contigs.fasta and all_contig_annotations.csv were analysed using dandelion^88^ singularity container (v0.5.4) against IMGT reference sequences^89^. Further pre-processing, including removing ambiguously annotated and extra contigs, was done using dandelion (v0.5.6). Only cells with single-pair heavy and light chains were used for downstream analysis of VDJ repertoire features. Somatic hypermutation rates were quantified using immcantation tools^90^ using default settings. Clone definition was performed using dandelion based on identical V-J gene usage, identical junctional sequence length and 85% junctional amino acid sequence similarity. Clone networks were computed using dandelion and visualized per individual. Clonotype distances were calculated as pairwise Levenshtein distances.

### Differential cell type abundance testing

Enrichment across cohorts or between vaccination timepoints for each of the identified cell type was computed using sccomp^91^. Briefly, the method employs a Bayesian model based on sum-constrained independent Beta-binomial distribution and Hamiltonian Monte Carlo sampling to calculate the probability of an effect across conditions on cell type proportions. The effect of cohort on the baseline differences in cell type proportions was modelled as *cell_count_* ∼ *Cohort* specifying a fold-change effect threshold of 0.1. Significant effects were defined based on the reported false discovery rate (FDR) being < 0.05 or < 0.1, respectively when comparing the differences across cohorts either for the broadly annotated or for the high-resolution cell types. The effect of vaccination on the differences in cell type proportions was modelled separately for each cohort as: *cell_count_* ∼ *Visit* + (1|*Donor ID*), specifying Donor ID as a random variable to account for repeated measurements and a fold-change effect threshold of 0.1. Significant differences between vaccination timepoints were defined as FDR < 0.1. The Seurat MixingMetric^92^ function with default parameters was used to quantify how well cells from different donors are mixed within high-resolution clustering of T cells. The resulting score ranging from 5-300 was then scaled to 1-0 using the min-max scaling approach defined as (*x* − min (*x*))/(max (*x*) − min (*x*)), with high values indicating poor donor mixing.

### Gene ontology enrichment and module expression

Gene ontology (GO) enrichment analyses were performed using clusterProfiler^93^. Genes associated with the gene ontology terms related to responses to LPS (GO:0032496), responses to type I, II and III interferon (GO:0034340, GO:0034341, GO:0034342, respectively) and with the canonical NF-kB signal transduction pathway (GO:0007249) were obtained from the GO.db R package (https://bioconductor.org/packages/release/data/annotation/html/GO.db.html). Average expression of the resulting set of genes across single-cells was obtained using the AddModuleScore function from Seurat.

### Differential gene expression analysis

Significant gene expression differences between cohorts at baseline were calculated using FindMarkers in Seurat, specifying a minimum absolute log fold-change threshold of ≥ 0.25 and an adjusted p-value threshold of ≤ 0.05. Batch-corrected log2 counts per million (CPM) were calculated from the aggregated pseudobulk counts and used to visualise the per-donor expression of each identified differentially expressed genes in the relevant cell type.

### Flow cytometry analysis

Cryopreserved PBMCs were thawed and stained with anti-CD19-APC (clone HIB19, BioLegend, San Diego, USA), anti-CD138-FITC (clone DL-101, BioLegend, San Diego, USA), anti-IgD-PerCP/Cy5.5 (clone 1A6-2, BioLegend, San Diego, USA), anti-CD27-BV605 (clone O323, BioLegend, San Diego, USA), anti-CD3-PerCP (clone UCHT1, BioLegend, San Diego, USA). After staining, cells were diluted in 2mL FACS buffer (PBS with 2% heat inactivated foetal bovine serum and 1mM EDTA) for immediate sorting. Single cells were analysed using the Cytek Aurora Cell Sorter at room temperature with 405 nm (45 mW), 488 nm (95 mW), 561 nm (45 mW) and 641 nm (55 mW) lasers. Gated singlet CD19^+^, CD138^-^ B cells were analysed for naïve B cells (IgD^+^, CD27^-^), unswitched memory B cells (IgD^+^, CD27^+^) and switched memory B cells (IgD^-^, CD27^+^) population frequencies.

### Whole genome sequencing and variant analysis

Genomic DNA (gDNA) for each patient with SAD or CVID was isolated from ∼1M PBMCs using the Quick-DNA Miniprep Plus kit (Zymo Research) following the manufacturer’s protocol. Purified gDNA was quantified and quality checked using Qubit BR dsDNA assay (Invitrogen) and via Nanodrop spectrophotometer. Libraries were prepared at the Australian Genomics Research Facility using Illumina® DNA PCR-free Prep kit and sequenced on the NovaSeq X Plus (150PE reads) to >30X for 18/20 libraries. FASTQ files were processed with Sarek (v3.5.1)^94^ using default settings to produce annotated germline variant calls. Briefly, after adapter trimming, reads were aligned to the GATK.GRCh38 reference genome with BWA-mem (v0.7.18)^95^ and processed with GATK (v4.5.0)^96^, including marking duplicate reads and error correction with BaseRecalibrator and ApplyBQSR. Germline SNPs and indels were called using HaplotypeCaller before quality filtering of variant calls with BCFtools (v1.2.0)^97^ and annotated by SnpEff (v5.1d, snpeff_db = GRCh38.105) and Ensembl VEP (v113.0, cache_version = 113)^98^. Coding variants within a gene set from the IUIS Inborn Errors of Immunity Committee (v2, July 2024, https://iuis.org/immunology-databases/) were filtered for depth (>10), allele frequency (<0.01%; GnomAD v4.1.0)^99^, ClinVar ^100^annotation (not benign) and Combined Annotation Dependent Depletion (CADD, v1.7.0)^101^ scores >20 before manual variant curation and assessment.

## DATA AVAILABILITY

Processed single-cell gene expression, cell surface protein and sgRNA count matrices are available at Zenodo (10.5281/zenodo.20266085). All other raw and processed sequencing datasets are awaiting upload to online data repositories and will be made available upon request.

## SUPPLEMENTARY MATERIALS

**Supplementary Figure 1.**
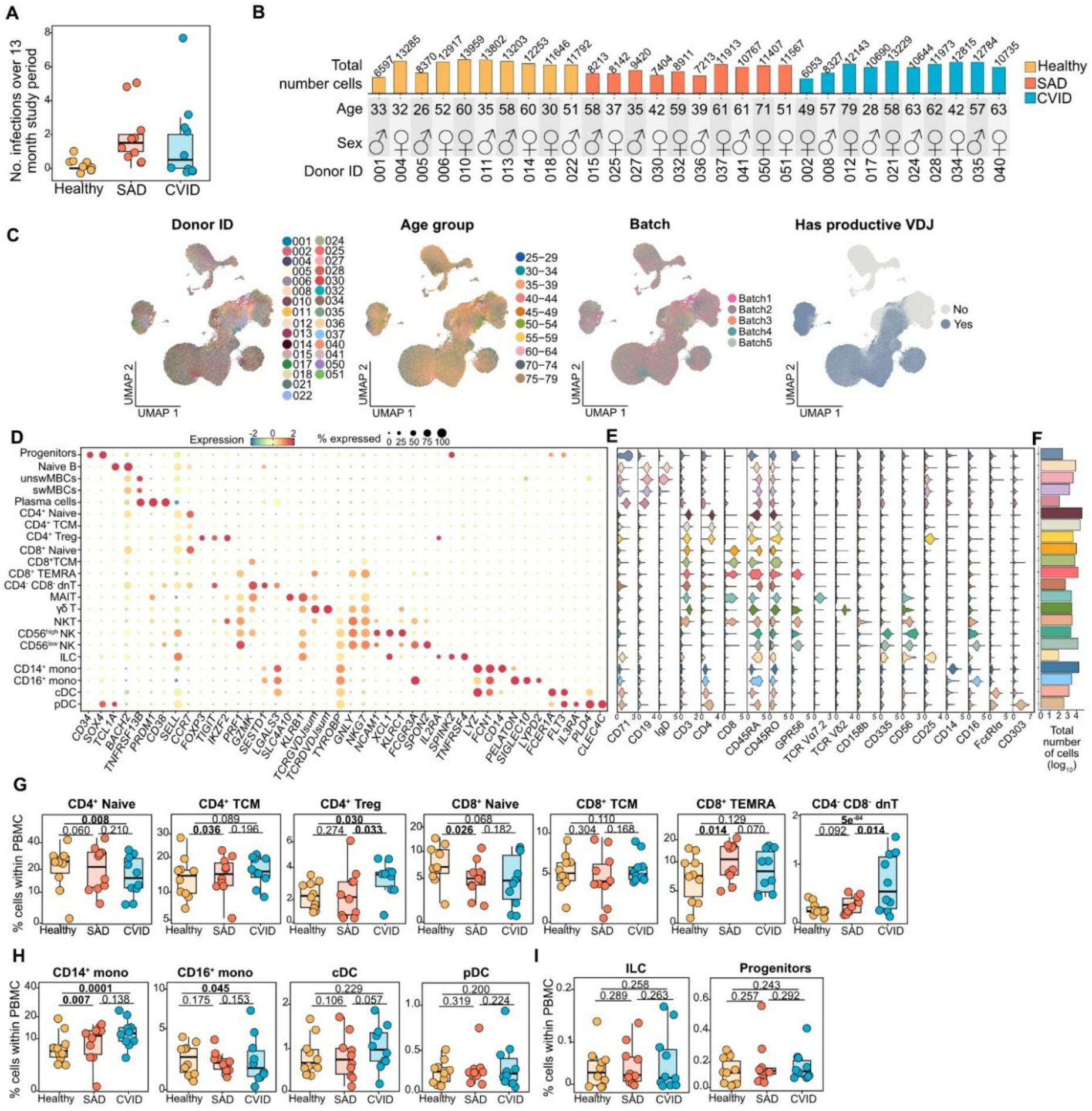
Overview single-cell atlas with cell type annotation and frequency across cohorts. **A)** Number of severe infections requiring hospitalization over the 13-month study period. **B)** Size of single-cell datasets per donor across cohorts, including donor age and sex. **C)** UMAP of single-cell CITE-seq resource by donor, age, batch, and productive VDJ data per cell. **D)** Expression of marker genes used for cell type annotation. Dot size depicts frequency of cells a gene is detected in. **E)** Expression of cell surface antibody-derived tag markers used for cell type annotation. **F)** Total number of cells per cell type. **G)** Relative frequency of T cell subsets, where *p* denotes the FDR-corrected probability of compositional effect between cohorts from sccomp. **H)** Relative frequency of myeloid cell subsets, as in G. **I)** Relative frequency of ILCs and progenitors, as in G.

**Supplementary Figure 2.**
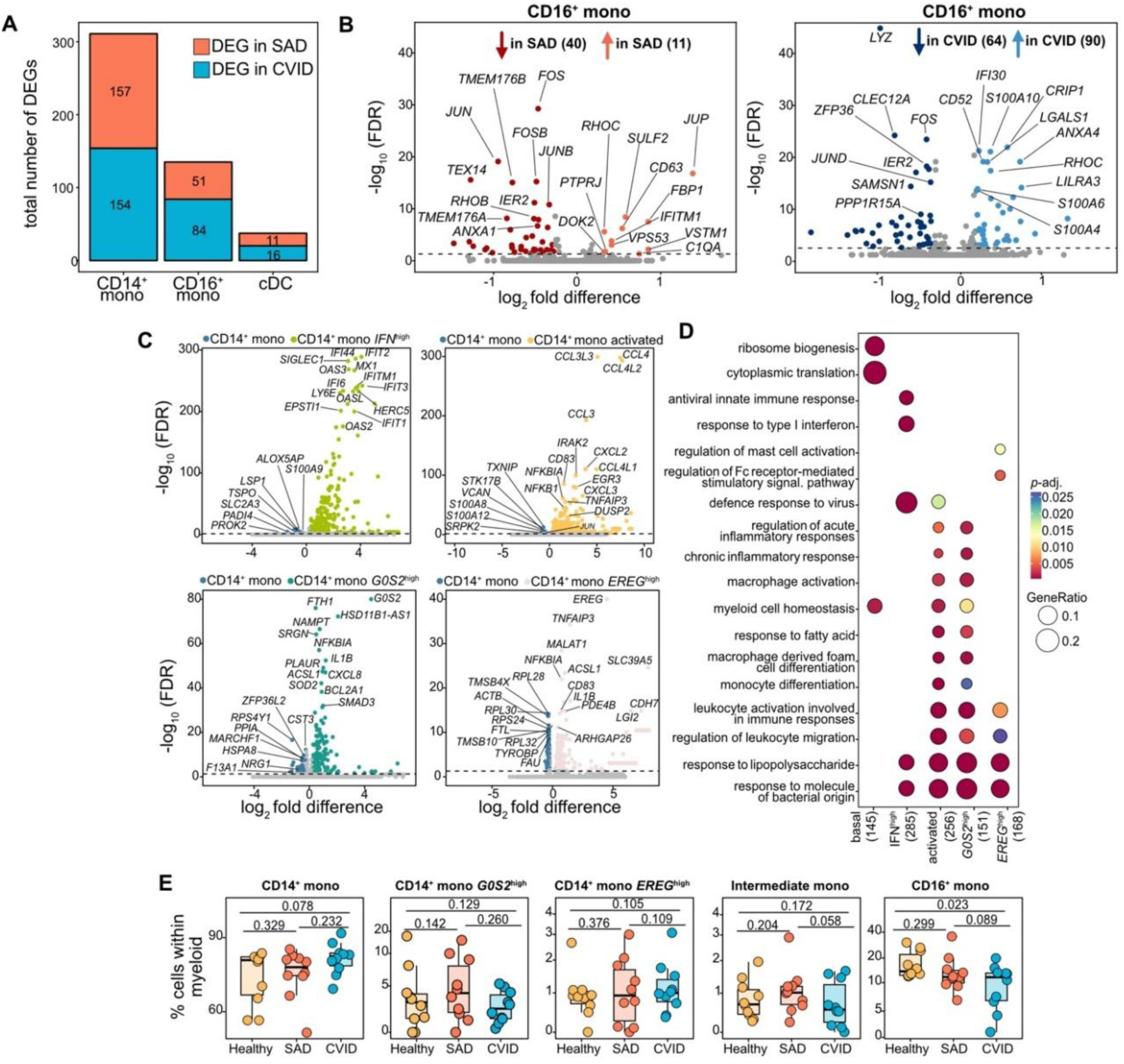
Variation in cell type frequency and gene expression across cohorts within monocytes. **A)** Number of differentially expressed genes between cohorts for CD14+ monocytes, CD16+ monocytes and conventional dendritic cells (cDC). **B)** Differential gene expression within CD16+ monocytes. **C)** Differential gene expression between subsets of CD14+ monocytes and basal CD14+ monocytes. **D)** Enriched gene ontologies for differentially expressed genes for the subsets of CD14+ monocytes. **E)** Relative frequency of monocyte subsets relative to total myeloid cells, where *p* denotes the FDR-corrected probability of compositional effect between cohorts from sccomp.

**Supplementary Figure 3.**
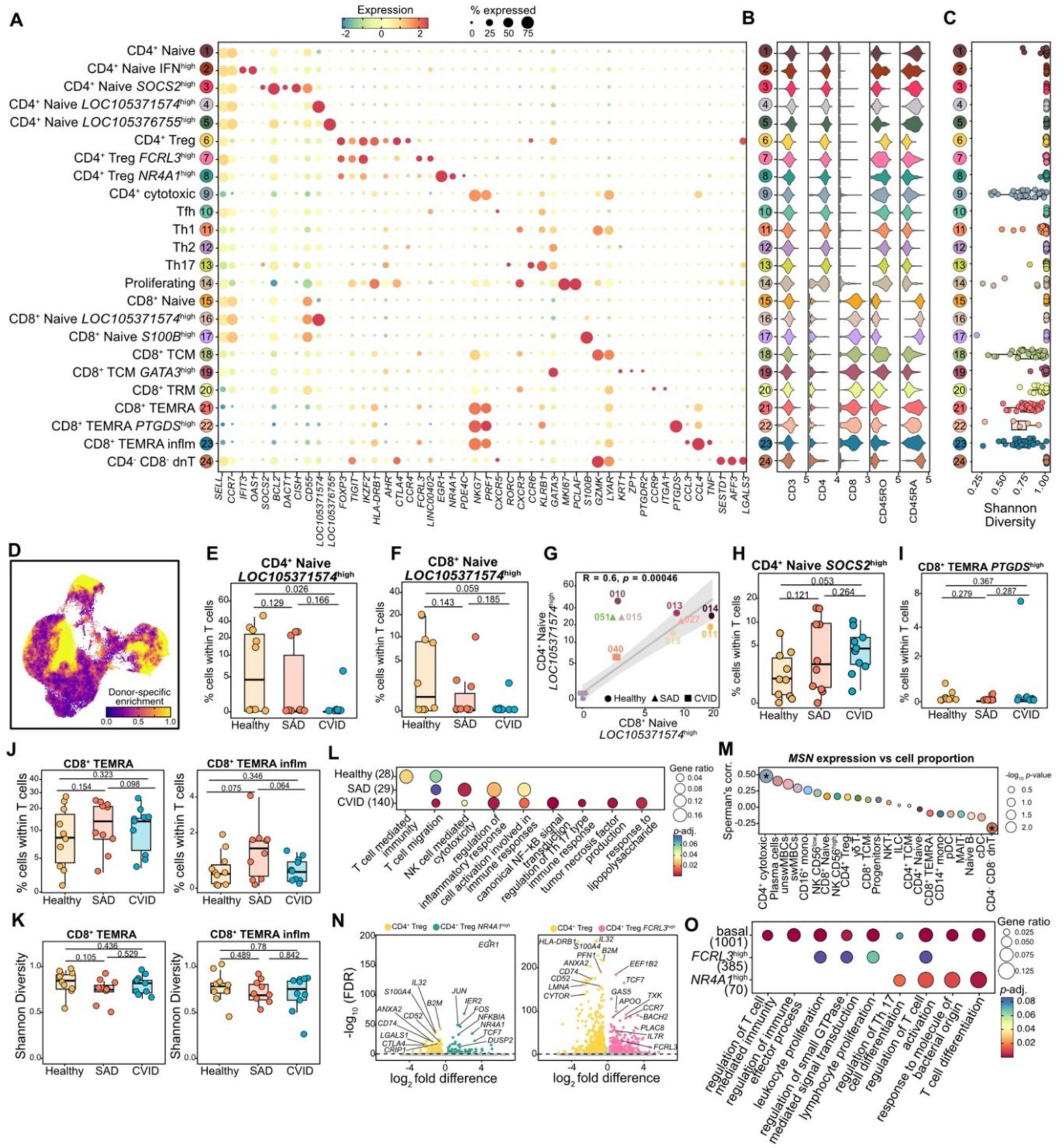
Overview of T cell annotation and variation in cell type frequency. **A)** Marker gene expression for high-resolution T cell subsets. Dot size depicts frequency of cells a gene is detected in. **B)** Expression of selected cell surface markers. **C)** Mean normalised Shannon diversity index across all single-cells for T cell subsets. **D)** Seurat MixingMetric between donors. High values indicate poor mixing across donors. **E-F)** Relative frequency of CD4+ naïve LOC105371574high (E) and CD8+ naïve LOC105371574high (F) relative to all T cells. *p* denotes the FDR-corrected probability of compositional effect between cohorts returned by sccomp. **G)** Correlation between CD4^+^ naïve LOC105371574^high^ and CD8^+^ naïve LOC105371574^high^ relative to all T cells across donors. **H-I)** Cell type frequency within T cells of CD4+ Naive *SOCS*high (H), CD8+ TEMRA *PTGDS*high (I). *P* denotes the FDR-corrected probability of compositional effect. **J)** Cell type frequency of CD8+ TEMRA and CD8+ TEMRA inflammatory relative to all T cells with *p*-values as in E. **K)** Mean normalized Shannon index of CD8+ TEMRA and CD8+ TEMRA inflammatory cells where *p* denotes Wilcoxon signed-rank sum test. **L)** Enriched gene ontologies for differentially expressed genes between cohorts for CD4+ cytotoxic T cells. **M)** *MSN* expression compared to cell proportion within PBMC across donors. Asterisks denote correlation *p*-values < 0.05. **N)** Differential gene expression between the subsets of regulatory T cells. **O)** Enriched gene ontologies for differentially expressed genes for the subsets of regulatory T cells.

**Supplementary Figure 4.**
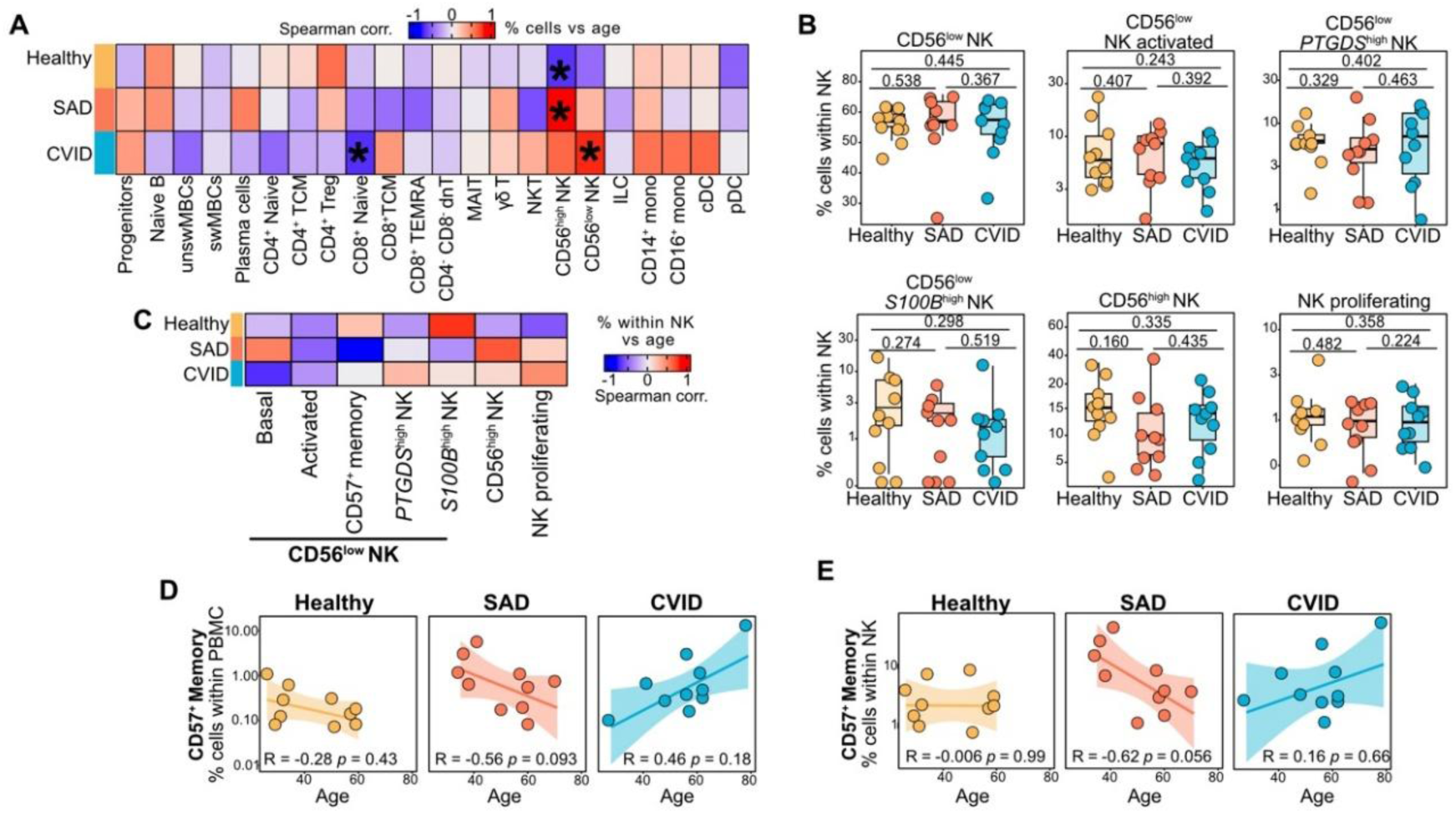
Association between NK cell type frequency differences with age. **A)** Association between individuals’ age and the frequency of each broadly annotated cell type within PBMC. Spearman’s correlation results shown. * denote *p*-values < 0.05. **B)** Relative frequency of the distinct NK subsets relative to total NK cells. *p* denotes the FDR-corrected probability of compositional effect between cohorts returned by sccomp. **C)** Association between individuals’ age and the frequency of each NK subset within NK cells. Spearman’s correlation results shown. * denote *p*-values < 0.05. **D)** Comparison between individual’s age and the frequency of CD56lowCD57+ NK memory cells relative to total PBMC. Spearman’s correlation results show. **E)** As in D but the frequency of CD56lowCD57+ NK memory cells is within NK cells.

**Supplementary Figure 5.**
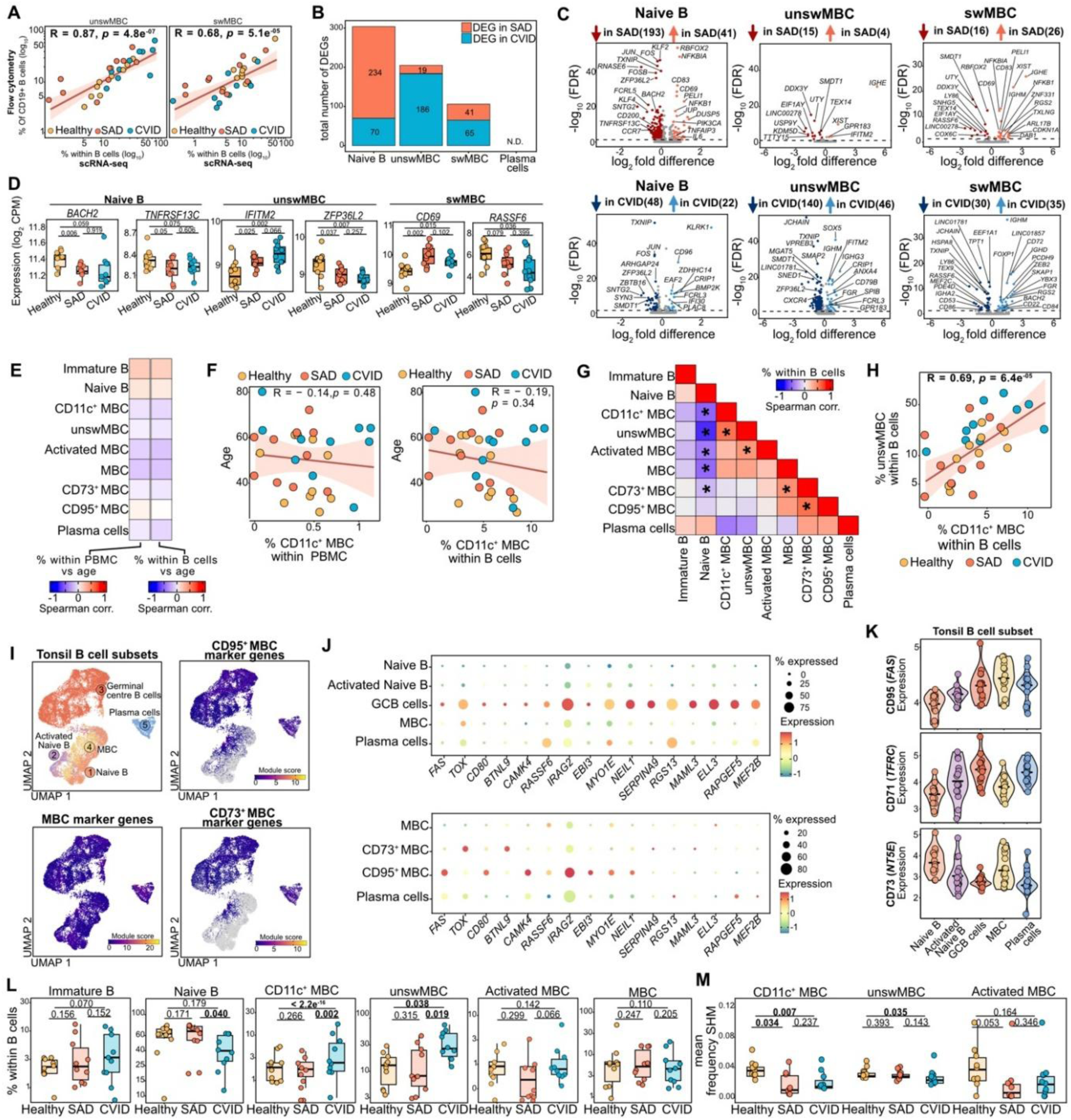
Gene expression and cell type frequency differences across cohorts within B cells. **A)** Comparison of different memory B cell (MBC) subsets defined by FACS or in the single-cell dataset. Spearman’s correlation results shown. **B)** Number of differentially expressed genes across cohorts for naïve B cells, unswitched MBCs and class-switched MBCs. **C)** Differential gene expression between cohorts for naïve B, unswMBC and swMBC. **D)** Mean expression for selected genes. *p* denotes results of Student’s t-test. **E)** Association between individuals’ age and B cell subset frequencies relative to total PBMC (left) or within B cells (right). Spearman’s correlation results shown. **F)** Relationship between individual’s age and frequency of CD11c+ MBC relative to total PBMC (left) or within B cells (right). Spearman’s correlation results shown. **G)** Correlation between B cell subset frequencies across donors. Spearman’s correlation results shown. * denote *p* values < 0.05. **H)** Relationship between the frequency of CD11c+ MBC and unswMBC within B cells across donors. Spearman’s correlation results shown. **I)** UMAP of tonsil B cell subsets obtained from49. Cells are colored by their annotation (top left) or by the expression of the top 100 gene markers for CD95+ MBC (top right), MBC (bottom left) and CD73+ MBC (bottom right). **J)** Marker gene expression for the subset of GCB cells across tonsil B cell subsets (top) or within peripheral memory B and plasma cells (bottom). Dot size depicts frequency of cells a gene is detected in. **K)** Expression of selected cell surface proteins across tonsil B cell subsets. **L)** Relative frequency of the distinct B cell subsets relative to total B cells. *p* denotes the FDR-corrected probability of compositional effect between cohorts returned by sccomp. **M)** Mean frequency of somatic hypermutation per each high-resolution B cells subtype. *p* denotes results of Wilcoxon signed-rank sum test.

**Supplementary Figure 6.**
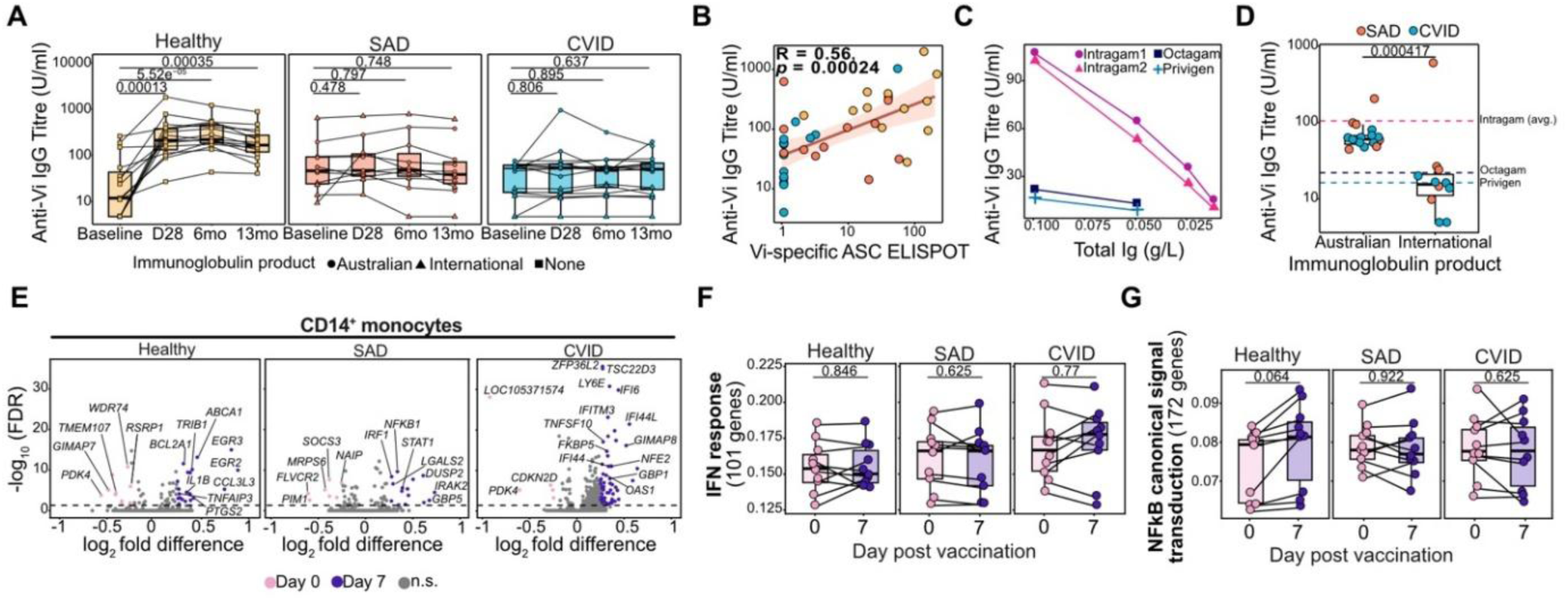
Differences in vaccination responses across cohorts. **A)** Quantification of anti-Vi IgG concentrations from serum samples collected for each individual across the 13-month study period. **B)** Spearman’s correlation of anti-Vi IgG titres measured at 28 days post vaccination with the frequency of Vi-specific ASCs at day 7. **C)**.Quantification of anti-Vi IgG concentrations in commercial immunoglobulin replacement therapy products (Intragam = Australian, Octagam/Privigen = International) **D)** Comparison of anti-Vi IgG titres in SAD and CVID individuals prior to vaccination (day 0) based on source of immunoglobulin replacement therapy product **E)** DEGs identified within CD14+ monocytes between baseline and 7 days post-vaccination for each cohort. **F)** Mean expression of 101 genes associated with IFN responses before and after vaccination within CD14+ monocytes. *p* values denote result of paired Wilcoxon signed-rank sum test. **G)** Mean expression of 172 genes associated with NF-kB canonical signal transduction before and after vaccination within CD14+ monocytes. *p* values denote result of paired Wilcoxon signed-rank sum test.

**Supplementary Figure 7.**
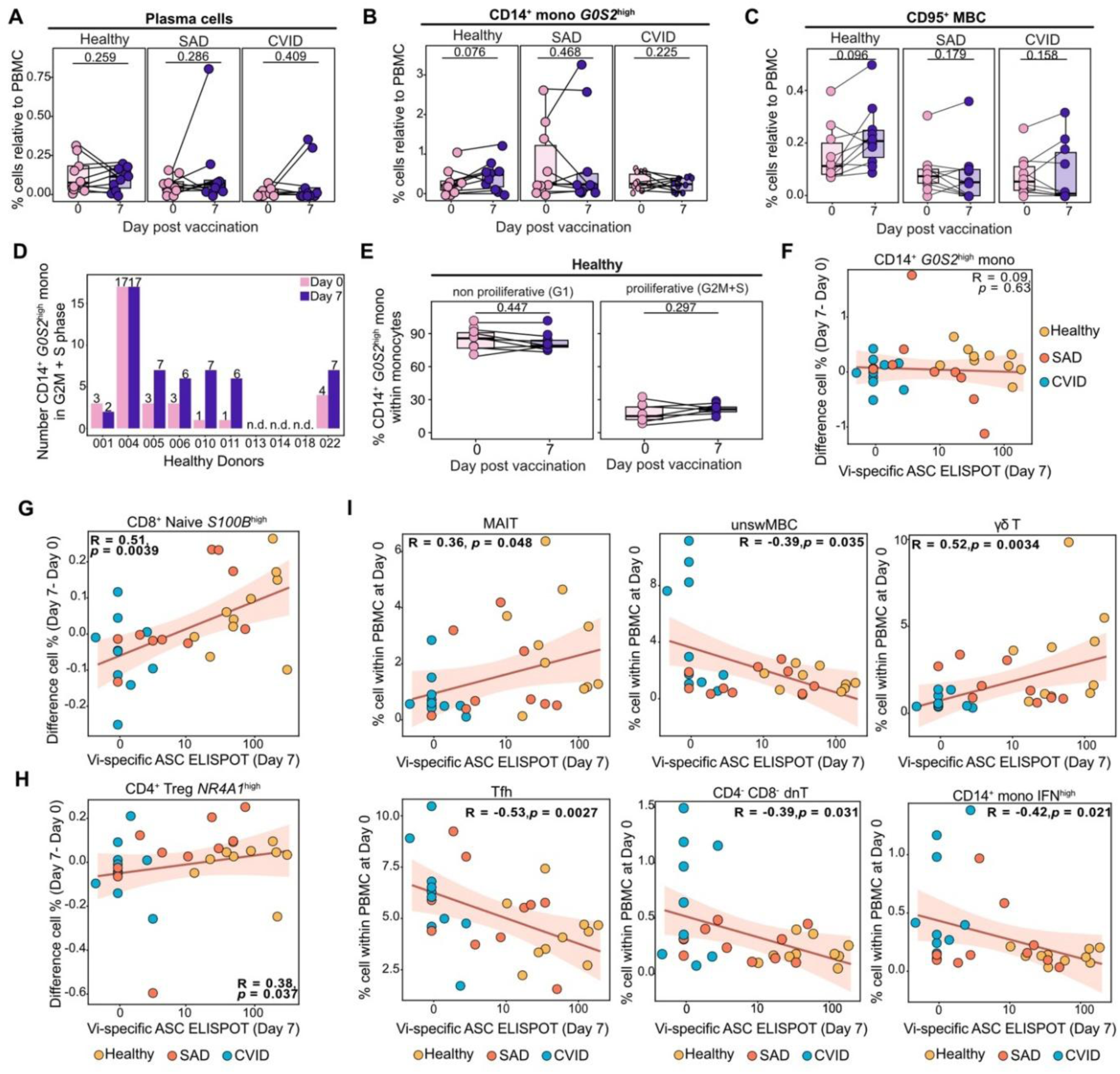
Correlates of vaccine responses. **A)** Proportion of plasma cells within PBMC before and after vaccination across cohorts. *p* denotes the FDR-corrected probability of compositional effect between cohorts returned by sccomp. **B)** Proportion of *G0S2*high CD14+ monocytes within PBMC before and after vaccination across cohorts. *p* as in A. **C)** Proportion of CD95+ MBC within PBMC before and after vaccination across cohorts. *p* as in A. **D)** Total number of cycling *G0S2*high CD14+ monocytes (G2M+S phase) before and after vaccination across healthy donors. **E)** Percentage of non-cycling (G1 phase) and cycling cells (G2M + S phase) within monocytes across controls between vaccination timepoints. *p* values denote result of paired Wilcoxon signed-rank sum test. **F)** Comparison of the difference in frequency of *G0S2*high CD14+ monocytes between vaccination timepoints with frequency of Vi-specific ASC (ELISPOT) at 7 days post-vaccination across individuals. Spearman’s correlation results shown. **G)** As F but for CD8+ naïve *S100B*high T cells. **H)** As F but for *NR4A1*high CD4+ regulatory T cells. **I)** Comparison of baseline frequencies of MAIT, unswMBC, γδ T, follicular T helper, CD4-CD8-dnT and IFNhigh CD14+ monocytes with the frequency of Vi-specific ASC (ELISPOT) at 7 days post-vaccination across individuals. Spearman’s correlation results shown.

Table S1. Donor information

Table S2. Raw counts broad resolution cell types at day 0 and day 7

Table S3. Raw counts high resolution cell types at day 0 and day 7

Table S4. Cell hashtag information

